# Mapping the global interactome of the ARF family reveals spatial organization in cellular signaling pathways

**DOI:** 10.1101/2023.03.01.530598

**Authors:** Laura Quirion, Amélie Robert, Jonathan Boulais, Shiying Huang, Gabriela Bernal Astrain, Regina Strakhova, Chang Hwa Jo, Yacine Kherdjemil, Denis Faubert, Marie-Pier Thibault, Marie Kmita, Jeremy M. Baskin, Anne-Claude Gingras, Matthew J. Smith, Jean-François Côté

**Affiliations:** Montreal Clinical Research Institute (IRCM), Montréal, QC, H2W 1R7, Canada; Molecular Biology Programs, Université de Montréal, Montréal, QC, H3T 1J4, Canada; Department of Chemistry and Chemical Biology and Weill Institute for Cell and Molecular Biology, Cornell University, Ithaca, NY, 14853, USA; Institute for Research in Immunology and Cancer, Université de Montréal, Montréal, QC, H3T 1J4, Canada; Department of Medicine, Université de Montréal, Montréal, QC, H3C 3J7, Canada; Department of Experimental Medicine, McGill University, Montréal, QC, H3G 2M1, Canada; Lunenfeld-Tanenbaum Research Institute, Sinai Health System, Toronto, ON, M5G 1X5, Canada; Department of Molecular Genetics, University of Toronto, Toronto, ON, M5S 1A8, Canada; Department of Anatomy and Cell Biology, McGill University, Montréal, QC, H3A 0C7, Canada

**Keywords:** ARF GTPases, BioID proteomics, ARF-Like GTPases (ARL), effector proteins, ESCPE-1, PLD1

## Abstract

The ADP-ribosylation factors (ARFs) and ARF-like (ARLs) GTPases serve as essential molecular switches governing a wide array of cellular processes. In this study, we utilized proximity-dependent biotin identification (BioID) to comprehensively map the interactome of 28 out of 29 ARF and ARL proteins in two cellular models. Through this approach, we identified ∼3000 high-confidence proximal interactors, enabling us to assign subcellular localizations to the family members. Notably, we uncovered previously undefined localizations for ARL4D and ARL10. Clustering analyses further exposed the distinctiveness of the interactors identified with these two GTPases. We also reveal that the expression of the understudied member ARL14 is confined to the stomach and intestines. We identified phospholipase D1 (PLD1) and the ESCPE-1 complex, more precisely SNX1, as proximity interactors. Functional assays demonstrated that ARL14 can activate PLD1 *in cellulo* and is involved in cargo trafficking via the ESCPE-1 complex. Overall, the BioID data generated in this study provide a valuable resource for dissecting the complexities of ARF and ARL spatial organization and signaling.

**SUMMARY STATEMENT:** Generation of the ARF family interactome allowed the attribution of potential localizations and functions to previously understudied members. We found that ARL14 activates PLD1 and contributes to ESCPE-1-mediated trafficking.

## INTRODUCTION

The ADP-ribosylation factors (ARF) family of small GTPases belongs to the RAS superfamily, together with RHO, RAS, RAN and RAB proteins. Members of the ARF family participate in a plethora of cellular processes including membrane trafficking, modulation of membrane lipid composition, and dynamic remodeling of the cytoskeleton (Sztul et al., 2019). Operating as GTPases, they are presumed to cycle between an inactive GDP-bound state and an active GTP-bound state. This cycling is orchestrated by guanine nucleotide exchange factors (GEFs), which promote GDP dissociation to allow GTP binding, and GTPase activating proteins (GAPs), which enhance the intrinsic GTP hydrolysis activity of ARFs (Sztul et al., 2019, Bourne et al., 1991). Unlike other RAS proteins that undergo lipidation at the C-terminus, the evolutionary conserved ARF family distinguishes itself due to a N-terminal extension. This extension forms an amphipathic helix in several family members, and is modified with various lipid groups (Gillingham and Munro, 2007). This helix plays crucial regulatory roles, being buried in a hydrophobic pocket in inactive ARFs and becoming exposed and deeply inserted into membranes concurrent with GTP loading (Amor et al., 1994, Antonny et al., 1997). This distinctive feature positions ARFs more proximal to membranes (1-2 nm) compared to other RAS superfamily proteins, creating a unique signaling environment (Gillingham and Munro, 2007). Since ARFs coupling to effectors occur on membranes of complex lipid compositions and of various curvatures, from flat to highly curved, and because of the challenges in capturing these events biochemically, our current understanding of ARF signaling remains fragmented (Sztul et al., 2019).

The ARF family consists of 29 members, categorized into 5 classical ARFs (Class I: ARF1, ARF3; Class II: ARF4, ARF5; and Class III: ARF6), 2 secretion-associated RAS (SAR1A and 1B) (Nachury et al., 2007), 21 ARF-likes (ARLs), and the tripartite motif-containing protein 23 (TRIM23) (Gillingham and Munro, 2007). Classical ARFs are primarily known for their role in recruiting coat proteins that initiate vesicle formation, facilitating membrane trafficking and cargo transport (Tan and Gleeson, 2019). For instance, ARF1, the founding member of the family, recruits the Coat Protein Complex I (COPI) at the Golgi membrane, enabling retrograde vesicular transport of cargos to the endoplasmic reticulum (ER) (Palmer et al., 1993). Additionally, ARF1 coordinates the formation of vesicles at the Golgi apparatus (hereafter referred to as Golgi) by recruiting the adaptor-proteins AP-1/3/4 (Dittie et al., 1996, Ooi et al., 1998, Hirst et al., 1999) and GGA proteins (Puertollano et al., 2001) for vesicular trafficking to endosomes or plasma membrane. The distinct functions of ARF3 and of ARF4/5 have been challenging to reveal since individual knockdowns do not induce major detectable phenotypes in cultured cells. However, their co-depletion with ARF1 has revealed specific Golgi functions, suggesting that they support ARF1 functions in the secretory pathway (Volpicelli-Daley et al., 2005). This underscores a high degree of interplay in ARF-regulated pathways. In contrast to ARF1, ARF6 predominantly operates at the cell periphery where it regulates processes such as endocytosis, endosomal recycling and cytoskeletal dynamics. This is achieved mainly through the modification of membrane lipid compositions via activation of PI(4,5)-kinase (Honda et al., 1999), and phospholipase D1 (PLD1) (Toda et al., 1999), as well as effects on actin through the activation of RHOA and RAC1 (Santy et al., 2005, D’Souza-Schorey and Chavrier, 2006).

Similarly to ARF1, SAR1A and SAR1B play crucial roles in ER-to-Golgi anterograde transport by recruiting the Coat Protein Complex II (COPII) and facilitating vesicle budding (Saito et al., 2017, Jones et al., 2003). As paralogues, SARs exhibit both overlapping and unique functions, attributed to differences in enzymatic and effector binding activities. For instance, SAR1B distinguishes itself from SAR1A with slower GTP exchange activity and higher affinity to SEC23 (Melville et al., 2020).

ARL proteins were initially identified by their sequence homology to ARF GTPases, yet they remain understudied in comparison to ARFs and SARs. ARLs control a wide array of biological functions across diverse cellular compartments (Kahn et al., 2006). Notably, ARL8A and ARL8B localize to lysosomes, facilitating their motility (Hofmann and Munro, 2006). Numerous other ARLs, including ARL2/3, ARL4D, ARL6, and ARL13B, play regulatory function in microtubule-associated processes within centrosomes, spindles, midbodies, basal bodies and cilia (Yong et al., 2020, Zhou et al., 2006, Bhattarai et al., 2019, Lin et al., 2020, Nachury et al., 2007, Li et al., 2010). The heterogeneous spatial distribution of some ARL proteins can drive interplay with multiple cellular processes. For example, ARL2 interacts with β-Tubulin and the Tubulin chaperone TBCD, promoting cytosolic biogenesis of α/β-Tubulin dimers (Francis et al., 2017a, Francis et al., 2017b). Additionally, ARL2 localizes to the intermembrane space of mitochondria, where it drives mitochondrial fusion (Newman et al., 2017). The intricate biological functions of individual ARL GTPases are complicated by their interplay with other ARFs. ARL1 crosstalks with ARF1/3, influencing the recruitment of the GEF BIG1 at sorting endosomes (D’Souza et al., 2014). Likewise, ARL13B acts as a GEF for ARL3, orchestrating the delivery of cargos into the cilium in cooperation with the ARF GAP

RP2 (Gotthardt et al., 2015, Veltel et al., 2008). Despite their importance, the effectors of ARLs are still largely unknown, highlighting the necessity of elucidating their cellular functions (Sztul et al., 2019). Moreover, there are still few data describing the biochemical properties underlying ARL nucleotide cycling or their propensity to function as archetypal “switch-like” GTPases. Many ARL proteins, including ARL5C, ARL9, ARL10, ARL11, ARL13A, ARL14 and ARL16, have yet to be assigned clear biological functions. Clearly, new approaches are needed to access unexplored layers of ARF and ARL spatial signaling, as well as to uncover the full repertoire of their associated proteins in cells.

Defective ARF/ARL signaling contributes to numerous human diseases. Many genetic diseases are linked to mutations in coat complexes and are collectively known as “Coatopathies”. These encompass mutations in COPI (Watkin et al., 2015, DiStasio et al., 2017, Izumi et al., 2016), AP (Montpetit et al., 2008, Nesbit et al., 2013, Assoum et al., 2016, Abou Jamra et al., 2011, Slabicki et al., 2010), COPII (Boyadjiev et al., 2006, Bianchi et al., 2009, Garbes et al., 2015) and Retromer complex associated proteins (Gustavsson et al., 2015, Damseh et al., 2015). Additionally, mutations in ARF GEFs, crucial in coat assembly, were identified (Sheen et al., 2004). In cancer, disruption of ARF-controlled membrane trafficking promotes cell invasion and migration (Casalou et al., 2020), while defective effector signaling can enhance oncogenic signals, such as ARF5/6-driven activation of PI3K/AKT (Nacke et al., 2021). ARFs are also hijacked by pathogens, including bacteria, that directly subvert ARFs or their cognate effectors (Jimenez et al., 2016). ARF6 has been identified as an important host factor for the cellular entry of the SARS-CoV-2 (Zhou et al., 2022, Mirabelli et al., 2022).

To gain fundamental knowledge into the ARF and ARL signaling pathways and their relevance to cell biology and human health and diseases, we performed proximity labeling (BioID) coupled with mass spectrometry on 28 constitutively active human ARF family members to define the comprehensive ARFome. Our approach reveals the broad landscape of ARF/ARL effectors and defines specificity for GAPs. In a complementary study, the proximity interactome of ARF/ARL proteins was investigated by short-term biotin labeling using miniTurboID (Li et al., 2022). Leveraging BioID’s ability to provide a 24-hour history of protein interactions, we were able to interrogate the localization of candidate proteins to define ARF/ARL functional localization. Finally, we discovered a potential coat complex recruited via the newly identified PLD1 activator, ARL14, on endosomes, facilitating retrograde traffic.

## RESULTS

### Development of a BioID pipeline to define the ARF family proximity interaction network

ARF and ARL GTPases are dynamic proteins that rely on a unique N-terminal region for membrane insertion, crucial for their functional roles. Therefore, a microenvironment closer to membranes, compared to other RAS and RHO proteins, is crucial for ARF/ARL signaling. Capturing many ARF-mediated interactions *in vitro* using classical biochemical approaches, in the absence of a phospholipid scaffold would be challenging. Instead, we elected to use BioID, a powerful method that has proven effective in systematically studying RHO GTPases in their native environment (Bagci et al., 2020). BioID utilizes an abortive biotin ligase derived from *Escherichia coli* that when fused to a protein of interest, labels proximal interactors with biotin within a 10 nm radius in the native micro-environment (Roux et al., 2012). Irreversibly biotin-tagged proteins are affinity-purified with streptavidin and identified by mass spectrometry at high throughput (Nahle et al., 2022). We systematically performed BioID following 24 hours of labeling time with 28 out of 29 ARF family members as baits. TRIM23 was not included in the study due to unsuccessful cell line generation. We chose to insert the BirA* tag at the C-terminal end of ARF/ARL proteins to leave the lipidated N-terminal extension unmodified (Fig. 1A).

**Figure 1.**
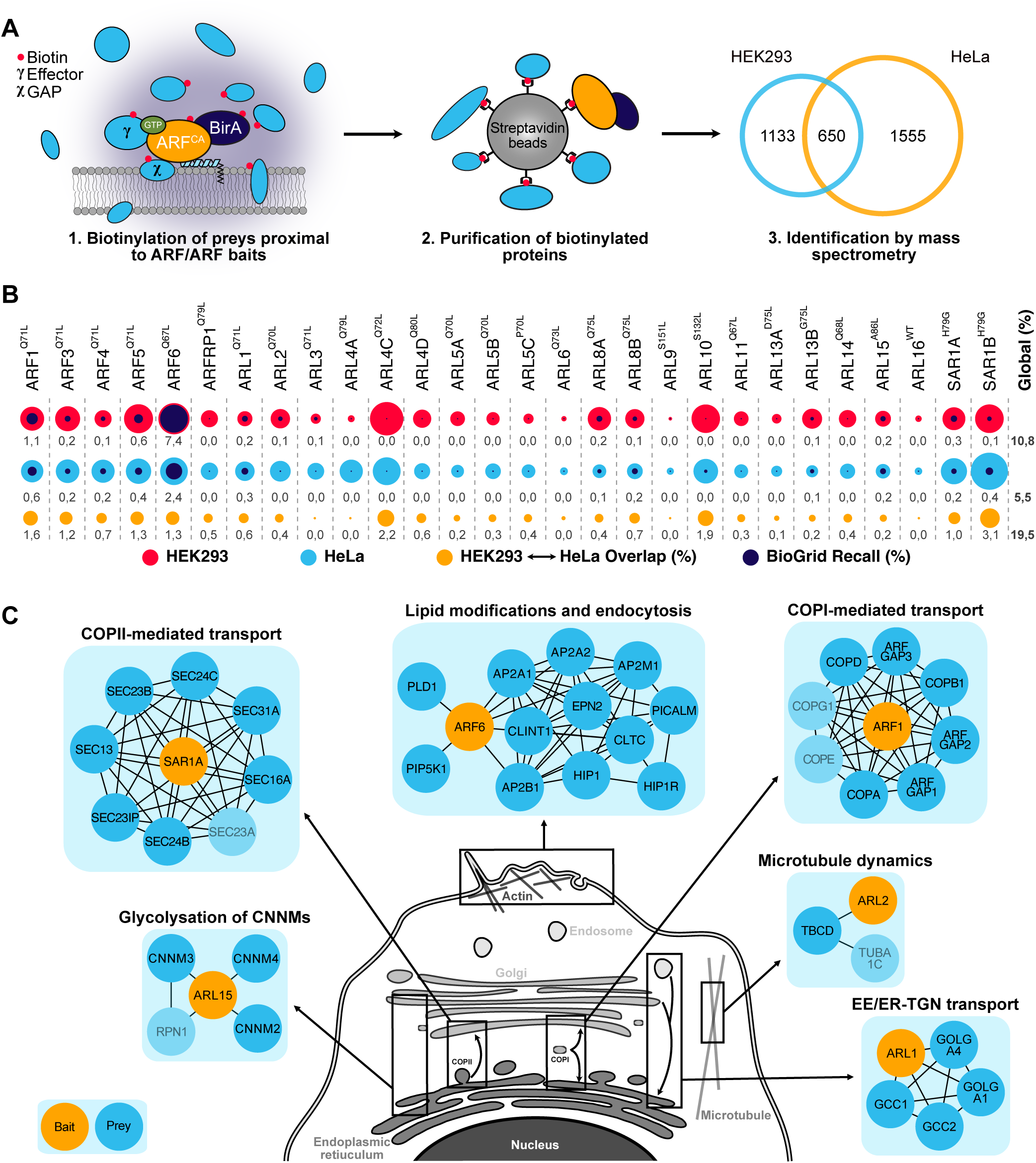
The ARF proximity interaction network defined by BioID. **(A)** Overview of the ARF BioID pipeline, performed in Flp-In T-REx HEK293 and Flp-In T-REx HeLa cell lines. The procedure involves inducing the expression of the constitutively active (CA) ARF family baits fused to BirA*-Flag with tetracycline for 24h in media supplemented with biotin (red dots). This permits the biotinylation of proximal proteins, including effectors (γ) and GAPs (χ) (Step 1). Subsequently, the biotinylated proteins are isolated (Step 2) and identified by mass spectrometry (Step 3). High confidence interactors were identified based on criteria: an AvgP ≥ 0.95, a CAAX-ratio of over 1.7, and spectral counts surpassing 4.5 in HEK293 cells and 6 in HeLa cells. The Venn diagram illustrates the number of interactions unique to each cell line, as well as those shared between them. **(B)** A proportional area chart illustrates the interactions identified for each bait (HEK293 in red and HeLa in light blue) and their overlap with existing literature (dark blue), proportionate to the total interactions in both cell lines. The orange circle represents the interaction overlap between HEK293 and HeLa cells, with the light grey numbers denoting the overlap percentage. The total overlap percentage and the total recall percentage between the two cells line are presented under the global column. **(C)** Known interactors of ARF and ARL GTPases involved in various cellular processes, identified by BioID thus validating the method. Interactors depicted in lighter blue were detected in our screens but did not pass our stringent thresholding. EE; early endosomes, ER; endoplasmic reticulum, TGN; *trans*-Golgi network.

To enrich for effectors and GAPs, as was previously demonstrated for RHO proteins (Bagci et al., 2020), we generated constitutively active mutant of each ARF family GTPase. These BirA*-tagged GTPases were kept in a constitutively active state by mutating the glutamine in the highly conserved G-3 motif (WDXGG**Q**), which prevents GTP hydrolysis. There are no *in vitro* biochemical data describing nucleotide cycling for most ARFs, but early work on ARF1 determined a Q71L mutation disrupts GTP hydrolysis in a manner similar to RAS (Kahn et al., 1995). However, in contrast to the classical RAS GTPases the mutation in ARF1 does not alter the rate of nucleotide release or relative affinity to GDP/GTP. This suggests the ‘activating’ mutation drives only a low-level increase in GTP-loading, yet this remains valuable for interactome mapping. A 24-hour labeling time is thus highly advantageous to identify a comprehensive set of binding partners. For GTPases lacking the conserved glutamine, we generated specific mutants based on the position where the glutamine would normally be found: ARL5C (P70L), ARL9 (S151L), ARL10 (S132L), ARL13A (D75L), ARL15 (A86L), and SAR1A and B (H79G). ARL16 was kept in its wild-type form since the G-3 motif involved in GTP hydrolysis is atypical (RELGGC) (Dewees et al., 2022) and its biochemical properties are unknown.

Because ARF nucleotide cycling and the impact of RAS^Q61L^-derived mutations are largely unstudied, we performed and compared BioID interactomes for ARF1, ARF4, ARF6 and SAR1A in both their wild-type and active states (Table S7). This revealed a significant increase in the abundance of known effectors and GAPs when using the constitutively active mutants. Conversely, GEFs demonstrate an enrichment when the wild-type forms are used as baits, consistent with the knowledge that they preferentially interact with the GDP-bound GTPases (Fig. S1). These results support the utilization of active ARF/ARL mutants for identification of ARF binding partners, though their behavior might differ from RAS and RHO GTPases.

To establish the experimental controls, we employed BirA*-Flag and BirA*-Flag-eGFP to selectively filter out proteins biotinylated at random. Likewise, BirA*-Flag-eGFP-CAAX served as a control targeted to membranes, which filters for membranous preys with less than a 1,7-fold enrichment of a bait in comparison to this control. All baits used in this study, along with their mutants, are listed in Table S1. To ensure broad coverage and confidence in the proximity interactomes, we conducted BioID in two inducible cell lines: Flp-In T-Rex HEK293 and Flp-In T-Rex HeLa. HEK293 cells have been a classical model for proteomics and HeLa cells for their oncogenic profile and for convenience of imaging. One advantage of using inducible cell lines is the ability to mitigate potential long-term cellular damage caused by the expression of the proteins of interest.

Upon generating the cell lines, we conducted validation of both the expression levels the biotinylation capacity of all the baits by western blotting (Fig. S2). Additionally, we assessed the subcellular localization of each bait by immunofluorescence (Fig. S3). Co-staining experiments involving well-characterized ARF family members were performed to ensure that the tagging did not affect their localizations. Our results confirmed that the localization of ARF1, ARF3, ARF4 and ARF5 at the Golgi apparatus (Fig. S4A-B), while ARL8A and ARL8B exhibited colocalization with lysosomes (Fig. S4C). Also, the ER membrane resident SAR1A and SAR1B displayed a network-like immunostaining pattern adjacent to KDEL immunostaining, marking these proteins as targeted to the ER lumen (Fig. S4D).

### Using BioID to systematically delineate the ARFome

We systematically conducted BioID screens for all active baits, identifying over 63,000 proximity interactions (Table S8). Following stringent filtering (see Methods), we uncovered 1265 high-confidence interactors in HEK293 cells and 1671 in HeLa cells, with 532 proximity interactors shared between the two cell lines (Fig. 1A). The non-shared interactions between the two cell lines may be due to the different protein expression profiles, stringent thresholding, and the variations in the sensitivity of the two mass spectrometers used for HEK293 and HeLa cells. The overlapping preys for each bait between the two cell lines, along with a comparison to known interactions from the literature, are depicted in Figure 1B. Overall, approximately 10% of our hits in HEK293 cells and 6% in HeLa cells overlap with existing literature (Fig. 1B).

The quality of our BioID screens was benchmarked by leveraging previously established interactions. Within these interactions, the ARF1 dataset featured GAPs (ARFGAP1, 2 and 3) and the COPI complex (α-, β-, γ-, δ-and ε-COP), the later required for retrograde trafficking from Golgi to ER (Fig. 1C). Additionally, the complete COPII complex (SEC16A, SEC23A, SEC23B, SEC24B, SEC24C, SEC31A, SEC23IP and SEC13), essential for anterograde trafficking between Golgi to ER, was identified within the SAR1A dataset (Fig. 1C). ARF6 BioID retrieved known interactors involved in lipid composition modulation (PLD1, PIP5K) and endocytosis (AP2A1, AP2A2, AP2B1, AP2M1, CLTC, CLINT1, EPN2, PICALM, HIP1, HIP1R) (Fig. 1C). Furthermore, with ARL1 as a bait, the four metazoan GRIP domain golgins (GOLGA1, GOLGA4, GCC1 and GCC2), which require ARL1 for their targeting to the Golgi (Setty et al., 2003, Panic et al., 2003) (Fig. 1C). ARL2 was found proximal to TBCD and β-tubulin, forming a trimer important for maintaining microtubule densities (Francis et al., 2017b) (Fig. 1C). ARL15 acts as a negative regulator of cellular Mg^2+^ transport by modulating CNNMs N-glycosylation, with our BioID hits including, CNNM2, CNNM3, CNNM4, and ribophorin I protein (RPN1) (Zolotarov et al., 2020) (Fig. 1C). The recall of these well-characterized interactions underscores the quality of the BioID screens, providing confidence in the potential to reveal new protein interactions and uncovering unsuspected functions for both extensively and less explored ARF/ARL proteins. The comprehensive BioID data are available for exploration by the research community at http://prohits-web.lunenfeld.ca.

### Mapping the functional localization of ARF and ARL proteins *in cellulo* using BioID

Accurate determination of the localizations and the intricate vesicular trafficking between intracellular compartments, regulated by classical ARFs and atypical ARLs, has proven challenging using biochemical approaches or epifluorescence microscopy. Using BioID with a 24-hour labeling window enables the capture of a global overview of the proximal protein interactions occurring at specific subcellular locations during dynamic cellular processes such as vesicular trafficking. Hence, mapping the spatial distribution of ARF/ARL interaction with their effectors and their regulators holds great potential in deciphering additional molecular functions of ARFs and ARLs within specific organelles. Here, leveraging our BioID data, our aim was to assign to each bait its main intracellular localizations by querying the Human Cell Map database, a BioID-based online tool that compared 192 subcellular markers to predict bait localization (Go et al., 2021). Using this resource, we systematically predicted the top three functional localizations of each ARF/ARL (Fig. 2A). From our BioID results, at least one previously reported localization for most well-studied baits was identified. As proof of concept, analyses of ARF1 BioID indicated a localization to the Golgi, corroborated by the presence of various golgin family members (GOLGA3, GOLGA5, GOLGB1). Additionally, the tool suggested localization of ARF1 to recycling endosomes, where VPS50, part of the EARP complex was identified (Stearns et al., 1990, Kondo et al., 2012). Our BioID data also reveal that SNARE proteins (VAMP3, VAMP8 and VTI1B) from the early endosome became proximal to ARF1 (Fig. 2A and B), a localization predicted by the Human Cell Map tool. However, BioID data were unable to predict the expected localization for ARFRP1, ARL4A and ARL11. Instead, ARFRP1 was predicted to localize at the ER-Golgi intermediate compartment (ERGIC), the peroxisome and the nucleus; ARL4A at the ER and mitochondria; and ARL11 at the ER and ERGIC. The recapitulation of previously reported localization for most well-studied bait validated the efficacy of this approach. Therefore, mapping the localizations of previously known and understudied ARF and ARL proteins could unveil new layers of information.

**Figure 2.**
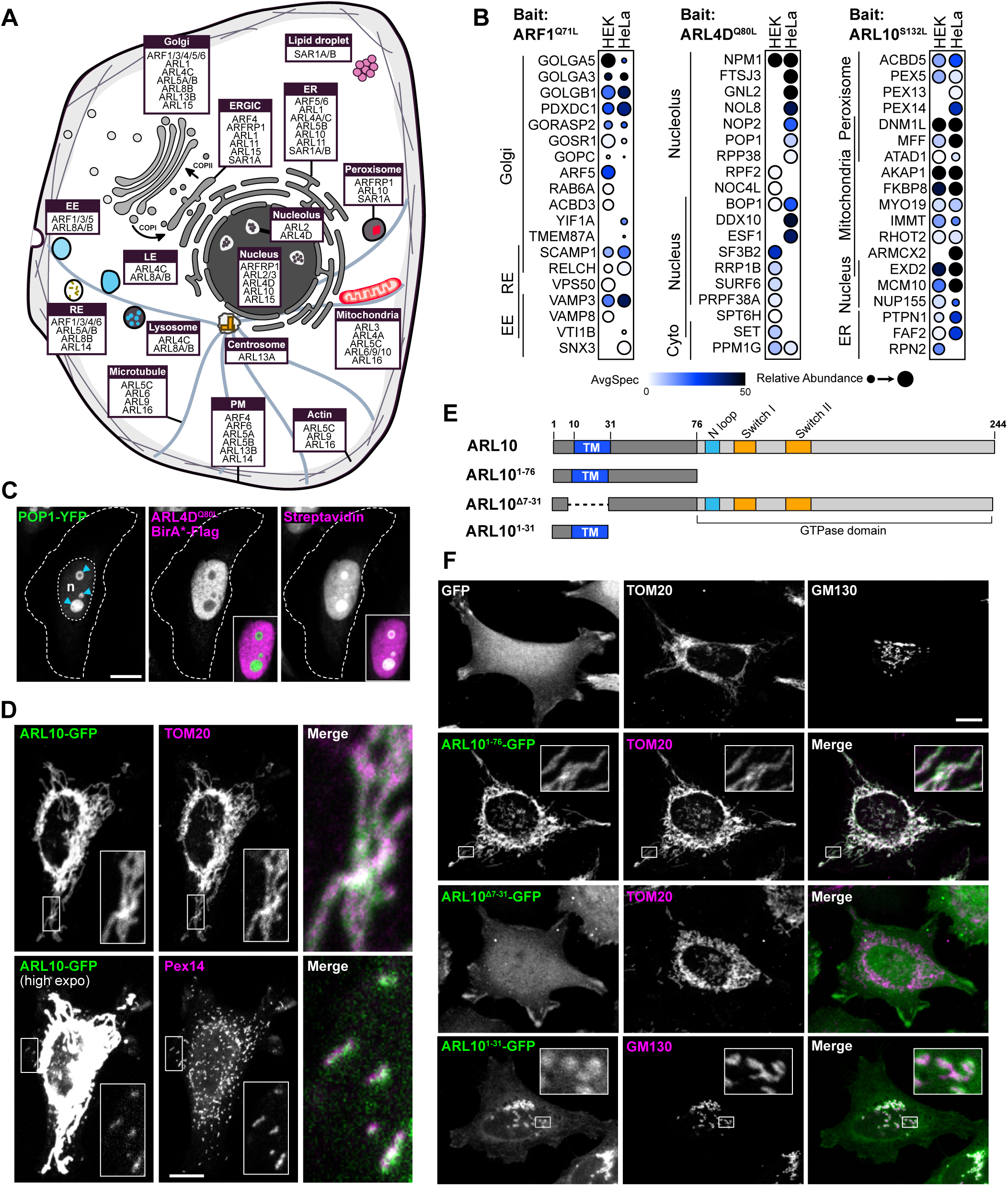
ARF location assignment from BioID results. **(A)** Graphical depiction of the top three cellular compartments enriched for each ARF/ARL as assigned by Human Cell Map tool following input of our BioID data. The top 4 is represented when a tie is present in the analysis. **(B)** Dotplots illustrating the prey enriched in the top 3 cellular compartments identified with Human Cell Map for constitutively active ARF1, ARL4D, and ARL10. Darker circles indicate higher spectral counts, while circle size represents relative abundance, AvgP ≥ 0.95. **(C)** FlpIn T-REx HeLa cells expressing POP1-YFP and constitutively active ARL4D-BirA*-Flag revealed with anti-Flag antibodies. Biotinylated proteins were detected using Alexa Fluor 633-conjugated streptavidin. Blue arrowheads indicate the nucleoli, and “n” denotes the nucleus. Insets presents a merge with POP1-YFP. Merges represent a ∼6.7x magnification. Bar, 10 µm. **(D)** Co-immunostaining of FlpIn T-Rex Hela expressing wild-type ARL10-GFP with anti-TOM20 (mitochondria marker) and anti-PEX14 (peroxisomes marker). Bar, 10 µm. **(E)** Schematic representations of ARL10 fragments used in this study. The N-terminal region is represented in dark gray and includes the putative transmembrane (TM) domain is represented in blue. The light gray portion represents the ARF GTPase domain, which include the N-loop, the Switch1, and the Switch2. **(F)** Immunostaining with anti-TOM20 and/or anti-GM130 (Golgi marker) was conducted on FlpIn T-Rex Hela expressing indicated GFP and ARL10 chimeric proteins, to define the mitochondria targeting signal of ARL10. Insets represent a ∼6.7x magnification. Bar, 10µm. EE: early endosomes, ER: endoplasmic reticulum, ERGIC: ER-Golgi intermediate compartment, LE: late endosomes, PM: plasma membrane, RE: recycling endosomes.

Leveraging the new information provided by the ARF family localization mapping described above, we examined deeper the functional distributions of the atypical ARL4D given that multiple subcellular compartments are reported in the existing literature. ARL4D undergoes myristoylation at its N-terminus, which is crucial for its membrane anchorage (Li et al., 2007), while also harboring a short basic extension at its C-terminus serving as a nuclear localization signal (Jacobs et al., 1999) (Fig. S5A). Consequently, previous studies documented the localization of ARLD at the plasma membrane, cytoplasm, and nucleus (Li et al., 2007). Using the BioID data on ARL4D, the Human Cell Map analyses indicated predominant localizations to the nucleus, cytoplasm, and nucleolus (Fig. 2A and B). Co-immunostaining of ARL4D with POP1, a nucleolus resident protein exhibiting proximity to ARL4D according to the BioID in HeLa cells (Fig. 2B), revealed exclusion of ARL4D from the nucleolus. However, biotinylated preys displayed pronounced colocalization with POP1 in the nucleolus (Fig. 2C). This result hints at a potential role for ARL4D at the interface between the nucleoplasm and the nucleolus, which would potentially be overlooked based strictly on ARL4D immunostaining.

Determining the nucleotide-loaded state of small GTPases in cells is extremely challenging and currently undetermined for all ARF family members, including ARL4D. To address this, we adopted an ion-pair reversed-phase high-performance liquid chromatography (IP-RP-HPLC) approach recently developed to resolve GDP/GTP loading of RAS proteins (Araki et al. 2021). The well-studied KRAS served as a control, and substantial GTP loading of the KRAS^Q61L^ mutant was detected following its purification from HEK 293T cells (59% GTP; Fig. S6A and B). Wild-type KRAS remained fully GDP loaded, as did wild-type ARF1 (Fig. S6C), consistent with early HPLC results obtained using bacterially-purified ARF1 (Randazzo and Kahn, 1994). Supporting the premise that an ARF1^Q71L^ mutant does not spontaneously load GTP (Kahn et al., 1995), this variant did not co-purify with any detectable nucleotide implying the mutation may fundamentally weaken nucleotide affinity. To resolve the nucleotide loaded in ARL4D, we purified both the wild-type and Q80L mutant with C-terminal EGFP tags from HEK 293T cells. Both wild-type ARL4D and the mutant variant were purified with no bound nucleotide (Fig. S6D). This is consistent with *in vitro* data showing ARL4A and ARL4C rapidly release nucleotide and are likely purified in the apo state (Jacobs et al., 1999), and explains why the Q80L mutant of ARL4D does not augment binding to some partners (Li et al., 2007). More data are required to fully understand nucleotide loading of ARFs and the impact of RAS-mimetic mutations, though ARFs have proven difficult to study *in vitro*.

For understudied ARLs, localization had not been previously assigned to ARL5C, ARL9, and ARL10. The BioID data suggests that ARL5C and ARL9 function within mitochondria, as well as in both actin and microtubule cytoskeletons (Fig. 2A). Human Cell Map predicts that ARL10 accumulates within mitochondria and peroxisomes, as well as in the endoplasmic reticulum and nucleus (Fig. 2A and B). Co-immunostaining of ARL10-GFP with the mitochondrial import receptor subunit TOM20 (TOM20) confirmed a significant mitochondrial localization of ARL10 (Fig. 2D). Moreover, high exposure of ARL10-GFP signals validated the presence of ARL10 in PEX14-positive peroxisomes (Fig. 2D), thus unveiling two new localizations for this unstudied GTPase. ARL10 lacks the classical N-terminal extension that is lipidated to favor membrane insertion of most ARF family members. Instead, ARL10 features a N-terminal sequence of 76 amino acids preceding the GTPase domain, with no homology to other ARF proteins (Fig. S5A). To investigate if this region encompasses the mitochondria-targeting signal of ARL10, we generated a chimera by fusing the amino acids 1-76 of ARL10 to GFP (ARL10^1-76^-GFP). The addition of ARL10^1-76^ proved sufficient to tether GFP to the mitochondria, as evidenced by its colocalization with TOM20 (Fig. 2E and F). Analysis of ARL10 N-terminal sequence using the membrane insertion prediction tool TMHMM suggested the presence of a putative transmembrane (TM) domain spanning amino acids 10-31. This region is primarily composed of hydrophobic residues and is followed by stretch of polybasic residues (PBR), a feature often observed in the N-termini of “signal-anchored mitochondrial outer membrane proteins” (Fig. S5A) (Walther and Rapaport, 2009). Deletion of the predicted transmembrane domain from the ARL10 sequence resulted in the loss of ARL10 mitochondrial localization (ARL10^Δ7-31^, Fig. 2E-F). However, a chimera where the first 31 amino acids of ARL10 are fused to GFP (ARL10^1-31^-GFP) was not sufficient to induce its translocation to the mitochondria (Fig. 2F). Instead, ARL10^1-31^-GFP accumulated in the Golgi apparatus, as evidenced by its partial colocalization with the Golgi marker GM130 (Fig. 2F). Taken together, these findings suggest that ARL10 is unique withing the ARF family by the presence of a TM domain, that cooperates with an adjacent PBR, for its efficient targeting to the outer membrane of the mitochondria.

### Mapping the ARF and ARL effectors and regulators

A major objective was to take advantage of the BioID data of constitutively active ARFs and ARLs to enable the identification of effector proteins. To determine the specificity of the effectors identified in individual datasets, we generated a WD summed (WDS) score computationally and conducted clustering to highlight the distinctiveness within the set of preys identified with a particular bait (Fig. 3A). The reliability of this approach is demonstrated by the clustering of baits with their closest relatives that share similar effectors. For instance, classical ARFs, except for ARF6, clustered together, as did ARL5A and B, ARL8A and B, and SAR1A and B (Fig. 3A). To further validate this method, we examined known unique interactions. For instance, the highest WDS score was associated with TBCD in the BioID of ARL2 in HeLa cells, and the highest WDS score in the ARL1 dataset was for the GRIP domain golgins (GOLGA1, GOLGA4) (Table S8), all of which are previously established interactions.

**Figure 3.**
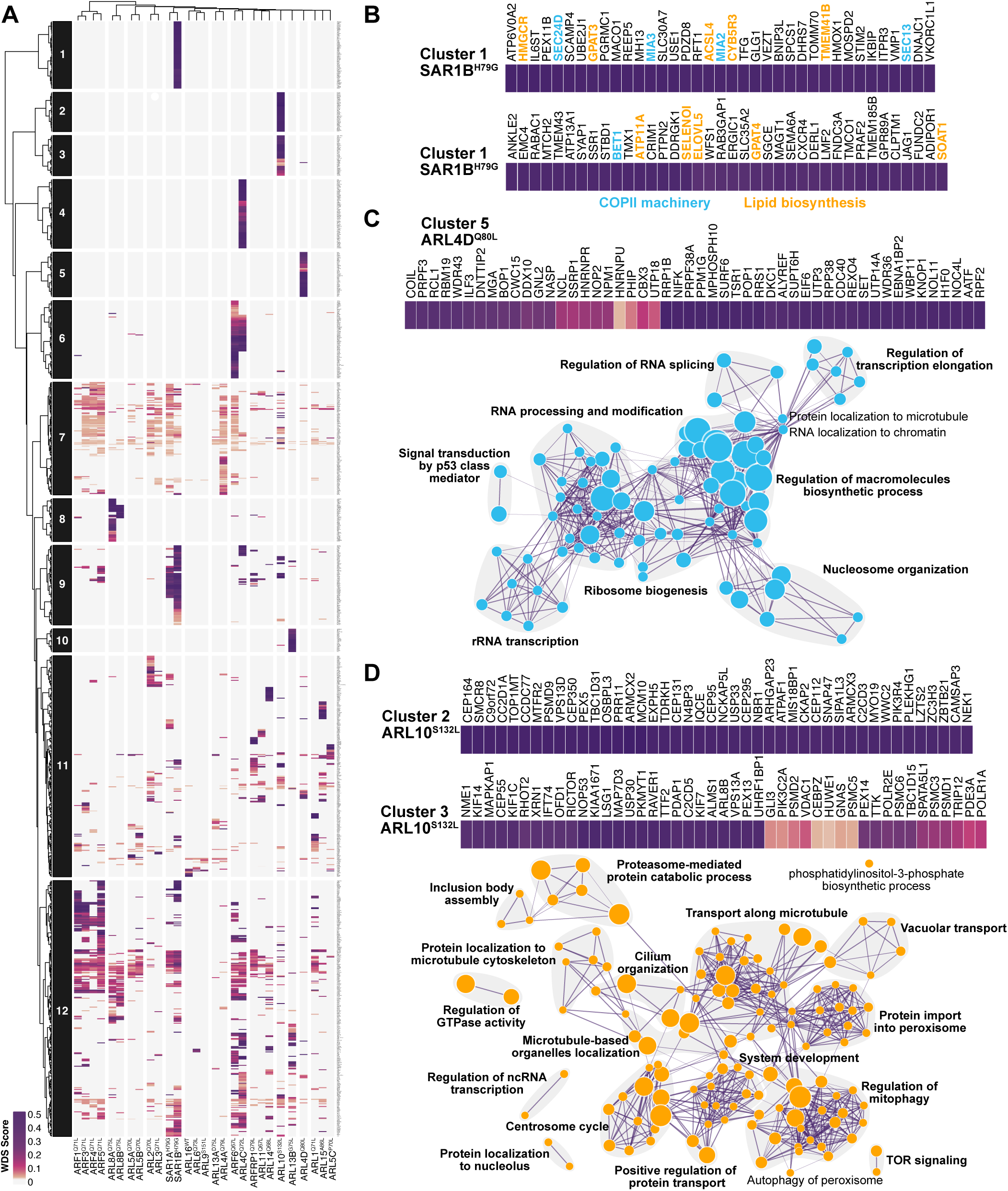
Clustering of ARFs and ARLs candidate effectors. **(A)** Heatmap of the specificity of potential effectors of ARF/ARL, determined by Pearson correlation and using the WDS score as metric. Dark purple signal represents the highest level of bait-prey specificity (highest WDS-score). **(B)** Zoom into Cluster 1, which is specific to constitutively active SAR1B, reveals the presence of proteins involved in the COPII machinery (light blue) and lipid biosynthesis (orange). **(C)** Zoom into Cluster 5, which is associated with constitutively active ARL4D, is shown at the top. The bottom panel represents an enrichment map of overrepresented Gene Ontology “Biological Processes” for this cluster. Node size reflects the number of proteins associated with each term, and the length of the edges indicates the interconnectivity between terms. Bold indicates grouped processes. **(D)** Zoom into Cluster 2 and Cluster 3, which are linked to constitutively active ARL10, is shown at the top. The bottom image presents an enrichment map of overrepresented GO “Biological Processes” of these clusters. Node size, edge lengths and lettering are as described in (C).

Clustering analyses revealed several distinct clusters, among them Cluster 1 was linked with SAR1B (Fig. 3B). SAR1B mutations, unlike SAR1A, are implicated in chylomicron retention disease, also known as Anderson disease, resulting in deficient chylomicron secretion and fat malabsorption (Jones et al., 2003, Shoulders et al., 2004). In Caco-2/15 cells, SAR1B silencing increased lipophagy, whereas SAR1A silencing resulted in the opposite phenotype (Sane et al., 2019), highlighting the functional differences between these SAR1 paralogs. These discrepancies are also mirrored by their differential proximity interactomes. Notably, components of the COPII machinery, such as SEC24D involved in cargo recognition and MIA2 involved in large cargo secretion from the ER like chylomicron (Santos et al., 2016), were uniquely associated with the SAR1B dataset. Additionally, certain preys exclusively passed our stringent BioID thresholding in SAR1B’s interactome, including BET1, which is a v-SNARE required for ER to Golgi transport (Mossessova et al., 2003) and TANGO1 (MIA3), which is involved in collagen secretion from the ER (Saito et al., 2011, Raote et al., 2020). Moreover, most of the proteins identified in Cluster 1 possess a transmembrane domain, with some involved in lipid metabolism (Fig. 3B). Together, these preys highlighted in the SAR1B dataset delineates avenues for further exploration in the machinery and cargos of SAR1B-associated COPII vesicles.

Pathway enrichment analyses of individual clusters revealed previously unknown localizations and functions for several ARLs. In particular, cluster 5 exhibits enrichment in preys specifically identified with ARL4D, recognized for its involvement in multiple cellular processes such as actin remodeling (Li et al., 2007), neurite formation (Yamauchi et al., 2009), adipogenesis (Yu et al., 2011), microtubule growth (Lin et al., 2020), and immune functions (Geers et al., 2022, Tolksdorf et al., 2018). Overrepresentation analyses of Gene Ontology “biological processes” are consistent with ARL4D’s residency in the nucleus, suggesting potential functions in the nucleolus. Additionally, roles in ribosome biogenesis, regulation of RNA splicing, regulation of transcription, nucleosome organization, regulation of macromolecules biosynthetic process, and RNA processing and modification are also indicated (Fig. 3C). Cluster 2 and Cluster 3 are uniquely associated with ARL10, an uncharacterized protein. Enrichment analysis suggests a role for ARL10 in regulating mitophagy, protein import into peroxisome, vacuolar transport, microtubule-based transport, cilium organization, centrosome cycle, and TOR signaling (Fig. 3D), consistent with its subcellular localization at the mitochondria and peroxisomes (Fig. 2D). To determine the nucleotide loaded state of ARL10 in cells we used the RP-IP-HPLC approach. Following purification of ARL10, the wild-type GTPase co-precipitated with both GDP and GTP, suggesting a significant fraction of ARL10 resides in an activated state (29% GTP; Fig. S6E). In contrast, the constitutively active (S132L) mutant of ARL10 did not bind detectible levels of GDP or GTP, indicating the mutation may reduce overall nucleotide affinity as observed with ARF1^Q71L^ and ARL4D^Q80L^. This result was not due to differential quantities of purified protein, as 4-fold more ARL10^S132L^ also lacked any detectable nucleotide (Fig. S6A and E). To further delineate ARL10 functional role and obtain insights into its protein-protein interactions, an affinity purification-mass spectrometry (AP-MS) experiment was performed on ARL10-GFP wild-type and the constitutively active mutant used in the BioID (Table S6). Most preys were identified with both ARL10 forms, albeit with variation in the number of spectral counts. Gene Ontology analysis of biological processes revealed enrichment in mitochondrion organization and transport, mitophagy and peroxisome organization for both variants. Additionally, the wild-type ARL10 displayed strong enrichment in processes related to organelle localization, organic acid metabolism, protein localization to peroxisome, and TORC2 signaling (Fig. S5B), suggesting the importance of nucleotide binding for these processes. The term “TORC2 signaling” was enriched in both the Cluster 3 from the BioID (Fig. 3D) and the AP-MS results, with core components RICTOR and MAPKAP1 representing mTORC2. The AP-MS also retrieved all other accessory proteins associated with this complex except for DEPTOR, highlighting the favorable interaction of wild-type ARL10 with these proteins (Fig. S5C). Co-immunoprecipitation confirmed this observation and the RP-IP-HPLC data, showing that ARL10 wild-type exhibits a stronger interaction with RICTOR than the constitutively active form (Fig. S5D).

We also found regulators of ARFs and ARLs nucleotide cycling activity in our BioID datasets. The constitutively active mutants selected for proteomics were predicted to associate preferentially with GAP proteins, being locked in GTP-loaded states. Interestingly, our data revealed an abundance of GAPs and GEFs in the datasets (Fig. 4). In line with previous findings, ARFGAP1/2/3 were proximal to the classical ARF1/2/4/5. While ARFGAP2/3 exhibited specificity, ARFGAP1 displayed a degree of promiscuity with several additional GTPases. AGFG1 was identified with class I classical ARFs (*i.e.* ARF1 and ARF3), class III (*i.e.* ARF6) as well as ARL8B and ARL13A. Furthermore, ARF6 was proximal to ACAP2, AGAP1 and AGAP3. In comparison to GAPs, the identified GEFs showed narrower specificity: ARFGEF2 was in the BioID of ARF1 and GBF1 was identified with ARL2. Additionally, GEF-like protein MON2 was identified with ARL1 and ARL10. Notably, ARL13B, a known GEF for ARL3, was identified in numerous ARF datasets. Collectively, our findings unveil several novel interactions between ARFs/ARLs and potential effectors, GEFs and GAPs. Future investigations should aim to further understand their biological significance.

**Figure 4.**
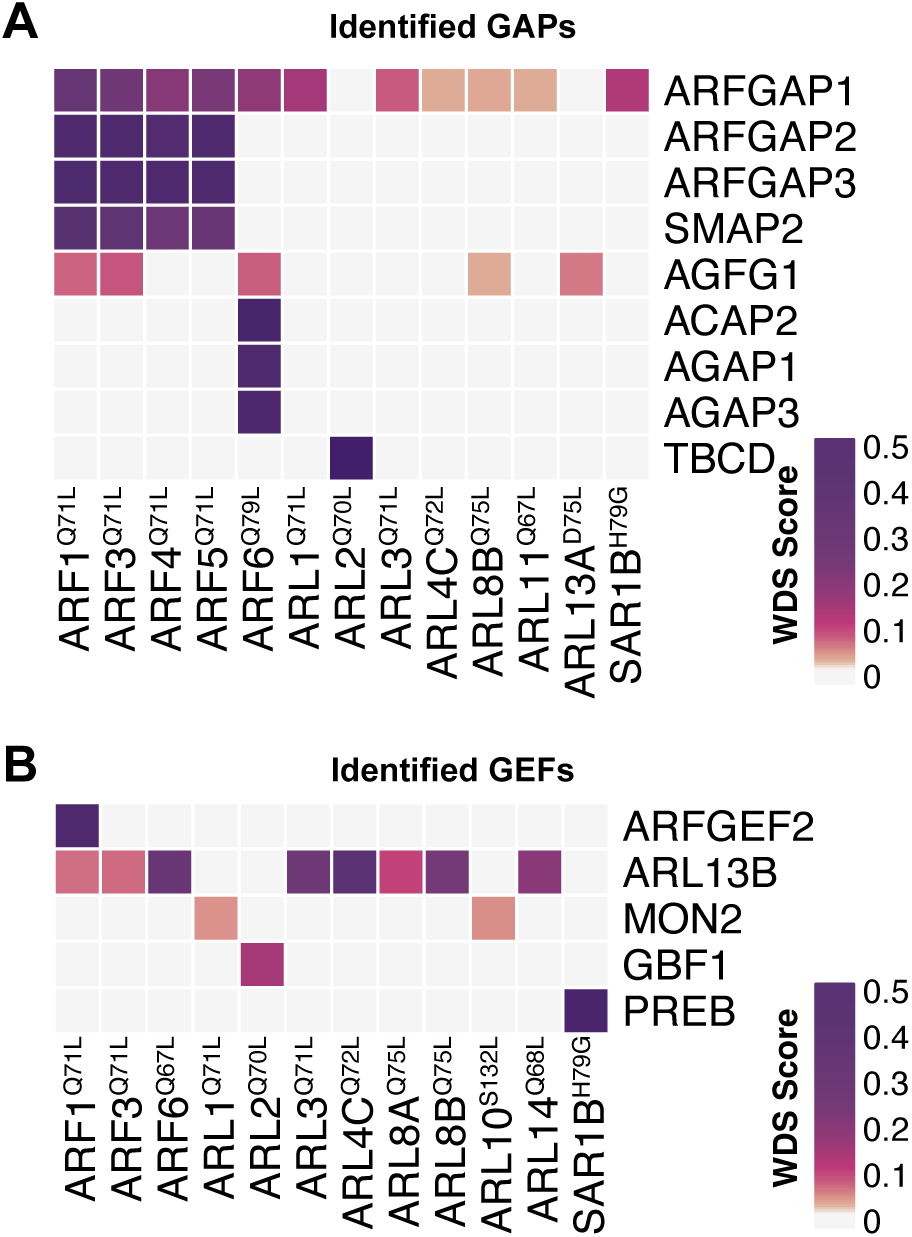
Mapping of ARF regulators. **(A and B)** Heatmap illustrates the specificity of GAPs in (A) and GEFs in (B), as identified through BioID. Dark purple indicates the highest level of bait-prey specificity, as determined by the WDS score.

### The understudied ARL14 is a robust activator of PLD1

Many members of the ARF family have remained relatively unexplored, with ARL14 backed only with a few reports for a role in lung cancer progression (Guo et al., 2019, Zhang et al., 2021). According to single cell mRNA expression data archived in the Human Protein Atlas, ARL14 expression is reported high in the gallbladder, stomach and intestines, with lower levels observed in the urinary bladder and pancreas (Uhlen et al., 2015) (proteinatlas.org). In the absence of suitable antibodies, we opted to validate the expression patterns of the ARL14 protein *in vivo* in mice by tagging the C-terminus of ARL14 with a 3xFlag epitope using the *i*-GONAD method (Gurumurthy et al., 2019). The mice were validated through genomic locus sequencing and two genotyping strategies (Fig. 5A and B). We extracted proteins from a range of organs from adult wild-type or transgenic ARL14^+/3xFlag^ mice, and western blotting analyses revealed exclusive expression of ARL14 in the stomach, small intestine, and large intestine (Fig. 5C).

**Figure 5.**
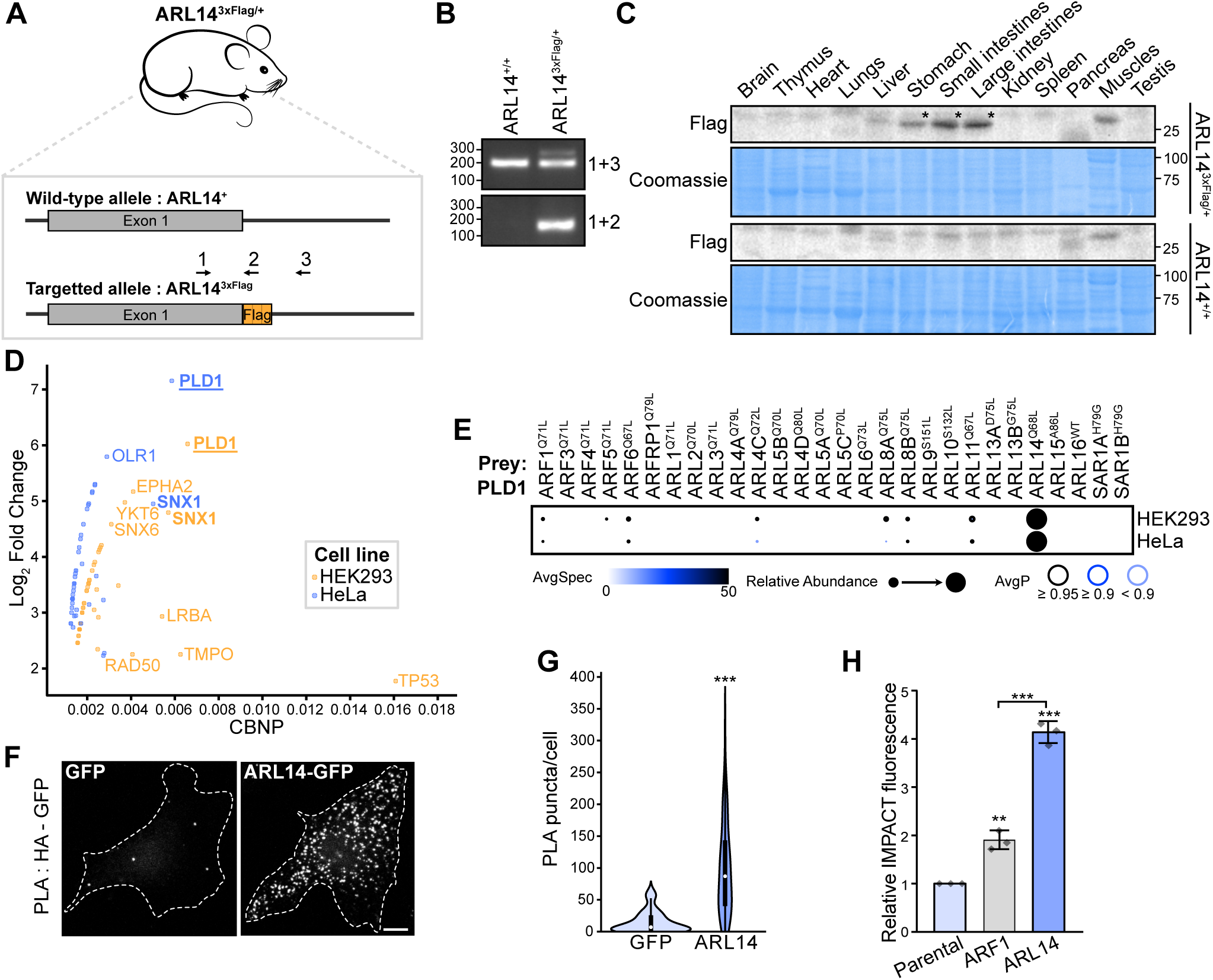
Characterization of ARL14, a PLD1 activator. **(A)** Schematic representation of the murine *ARL14* coding region, which consists of only one exon, targeted for genome editing to insert a 3x-Flag tag to the 3’ end. The genotyping strategy is indicated with arrows representing the different primers. **(B)** Genotyping results of mice generated by the strategy described in (A). Top represents PCR amplifications surrounding the insertion site (primer 1 and 3), while bottom represents similar PCR amplifications but with one primer specific for the inserted sequence (primer 2). **(C)** Western blots on proteins isolated from organs of ARL14^+/+^ (WT) or ARL14^3xFlag/+^ knock-in mice at age P65. The asterisk (*) indicates the bands of interest. Coomassie staining was used to assess loading of proteins. **(D)** Graph represents preys identified in ARL14 BioID, in HEK293 and HeLa cells, showing the normalized spectral counts (CBNP) on the X axis and the Log2 fold change against the mean of the negative controls on the Y axis. **(E)** A dotplot of the BioID results demonstrating proximity interactions of PLD1 with ARF GTPases in FlpIn T-REx HEK293 and HeLa cells. Darker circle color represents higher spectral counts, circle size denotes relative abundance, and circle outline corresponds to the indicated AvgP. **(F)** Proximity Ligation Assay (PLA) on FlpIn T-REx HeLa cells expressing HA-PLD1 and ARL14-GFP. Cells expressing HA-PLD1 and GFP alone were used as a control. F-actin staining (Alexa Fluor 568-Phalloidin) (not shown) was used to delineate the cell outline. Bar, 10 µm. **(G)** Quantitative analysis of (F) is presented, where the violin plot illustrates the number of PLA puncta per cell for each condition. Data are representative of n=3 independent experiments (total of 75 cells per condition). Statistics show the comparison of the GFP control to ARL14. *P* value was calculated by Student *t*-test; \*\*\**P*< 0.0001. **(H)** Measurements of PLD activity using IMPACT on active forms of the indicated GTPases, where the graph shows the relative IMPACT fluorescence quantified by flow cytometry. The bar height represents the mean, and error bars denote the standard deviation, n=3 independent experiments. Statistics above the graph are compared to the parental control. *P* values were calculated by one-way ANOVA, followed by Bonferroni’s test; \*\*\**P* < 0.0001, \*\**P*<0.001.

There are no biochemical data describing ARL14 function. To understand its nucleotide cycling, we first used IP-RP-HPLC of the constitutively active (Q68L) mutant and wild-type variants, which revealed no discernable nucleotide binding (Fig. S6F). As comparatively little ARL14 could be expressed and purified on beads (Fig. S6A), we attempted to study its cycling *in vitro*. Bacterially purified ARL14 was noticeably unstable and ^1^H-^15^N HSQC NMR spectra presented few robust or well-dispersed peaks characteristic of folded, nucleotide-bound small GTPases (Smith and Ikura, 2014, Killoran and Smith, 2019). This could not be improved by addition of GDP or a non-hydrolyzable analog of GTP (GTPγS; Fig. 6A). We corroborated these results by measuring the affinity of ARL14 to GDP and GTPγS using isothermal titration calorimetry (ITC), which confirmed ARL14 is unable to bind either nucleotide (Fig. 6B). These data validate the RP-IP-HPLC assay and suggest ARL14 is unlikely to bind guanine nucleotides. We considered if this was a general property of the subfamily which includes ARL4A/C/D and the closest relative, ARL11 (Colicelli, 2004). Purified ^15^N-ARL11 also presented poorly ordered resonances in HSQC spectra, but in contrast to ARL14 there was a significant improvement in chemical shift dispersion and lineshape upon addition of GTPγS, and to a lesser extent GDP (Fig. 6C). ITC experiments validated binding of ARL11 to GTPγS (Kd=1.53 nM) and GDP (Kd=1730 nM), corroborating the NMR results and establishing that ARL11 has an approximately 1000-fold higher affinity for GTP over GDP (Fig. 6D). This is a major deviation from classical RAS GTPases which maintain equal affinity to the two nucleotides, demonstrating significant biochemical divergence of some ARF GTPases that will undoubtedly impact function.

**Figure 6.**
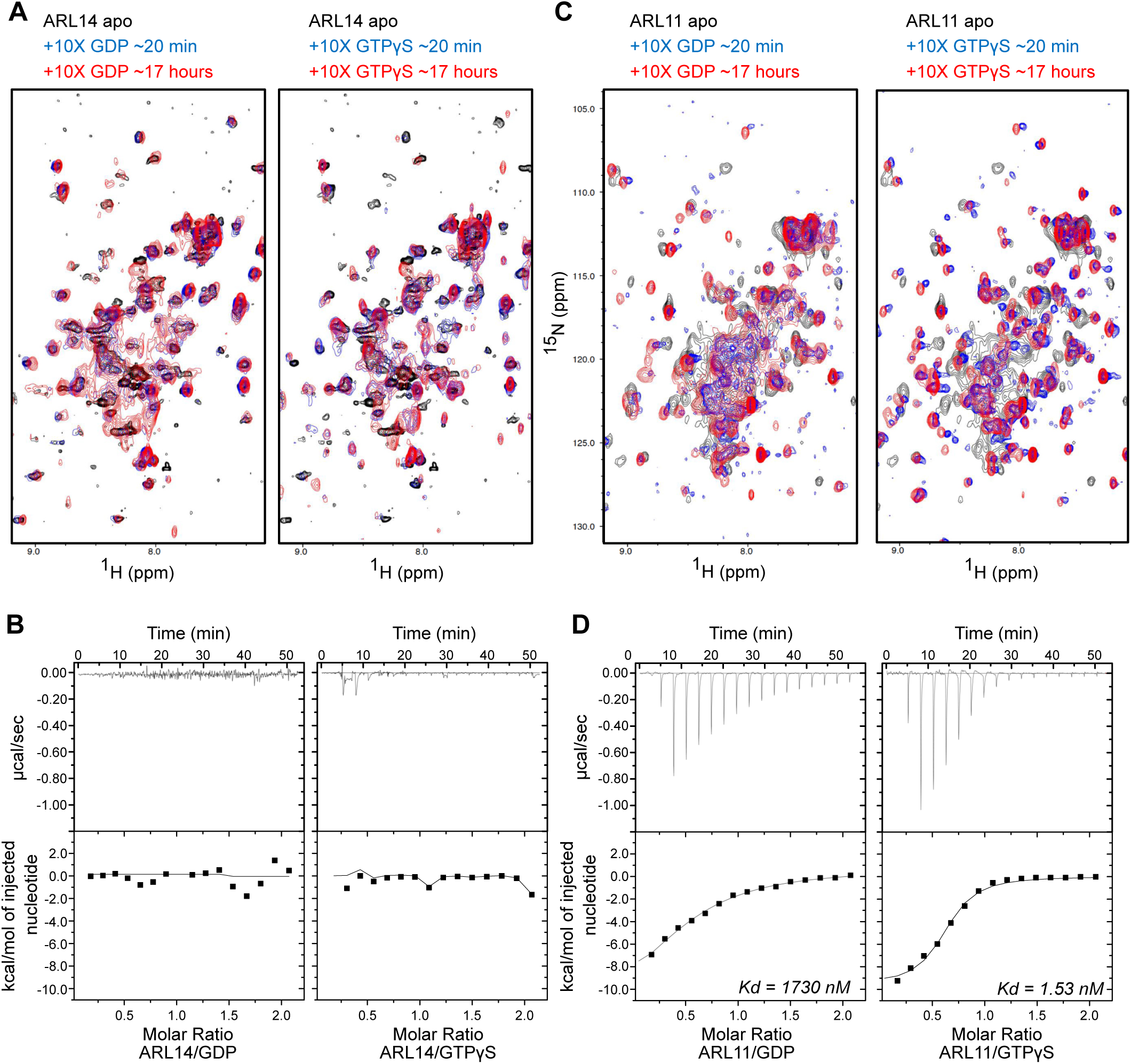
ARL14 has poor affinity for nucleotides. **(A)** NMR spectrum of purified ARL14 incubated with GDP or GTPγS. **(B)** Isothermal titration calorimetry analysis of ARL14 with GDP or GTPγS. **(C)** NMR spectrum of purified ARL11 incubated with GDP or GTPγS. **(D)** Isothermal titration calorimetry analysis of ARL11 with GDP or GTPγS. Kd; dissociation constant.

To investigate the molecular basis of ARL14 signaling, we opted to conduct experiments using only the wild-type variant to accurately reflect its nucleotide cycling *in cellulo*. We analyzed our BioID data to identify potential interacting partners. Among the notable proteins identified in the ARL14 interactome was PLD1. Our data exhibited significant spectral counts and the highest Log2 fold change compared to the mean of the negative controls for PLD1 in both HEK293 and HeLa cells (Fig. 5D). PLD1 functions by catalyzing the hydrolysis of its substrate, phosphatidylcholine, a rich phospholipid found in cellular membranes, yielding choline and phosphatidic acid (PA) products. PA, characterized by its cone-shaped structure, is a signaling lipid that facilitates negative curvature of membranes and also interacts with PA-binding proteins required for membrane fusion and fission (Zhukovsky et al., 2019, Roth, 1999). Notably, classical interactors of PLD1 include ARF1 and ARF6, which act as activators, establishing PLD1as an effector of these GTPases (Hammond et al., 1995, Frohman and Morris, 1999). While interactions between ARF1 and ARF6 and PLD1 were detected by BioID, the significantly higher PLD1 spectral counts observed in the ARL14 BioID experiments suggest ARL14’s potential role in activating PLD1 (Fig. 5E). Among the 28 ARFs/ARLs tested, low spectral counts for PLD1 were observed in the BioID studies of ARL4C, ARL8A and B, and ARL11, similar to those in the ARF1 and ARF6 experiments, implying a robust interaction with ARL14. To validate this interaction, we conducted a proximity ligation assay (PLA) in HeLa cells overexpressing ARL14-GFP and HA-PLD1, which confirmed that PLD1 is proximal to wild-type ARL14 (Fig. 5F and G). To assess whether ARL14 could activate PLD1, we used a click chemistry method named IMPACT, which stands for “Imaging PLD Activity with Clickable Alcohols via Transphosphatidylation” (Bumpus and Baskin, 2017, Bumpus et al., 2020). IMPACT involves two steps: (1) production of a clickable lipid via PLD-mediated transphosphatidylation of phosphatidylcholine with an exogenous alcohol (3-azidopropanol), and (2) tagging the clickable lipid with a fluorophore (BCN-BODIPY) via a click chemistry reaction. This experiment was performed in HeLa cells that inducibly overexpress either mScarlet, ARF1^WT^-mScarlet, or ARL14^WT^-mScarlet, and the fluorescent lipid product of the IMPACT reaction was quantified by flow cytometry, after thresholding on cells with similar mScarlet expression (Fig. 5H). This *in cellulo* approach demonstrated that ARL14 was a stronger PLD1 activator than ARF1, with a 4.5-fold and 2-fold increase, respectively, compared to the mScarlet control. These findings reveal ARL14 as a robust activator of PLD1.

### ARL14 is involved in ESCPE-1 mediated trafficking

In addition to PLD1, the <u>E</u>ndosomal SNX-BAR <u>s</u>orting <u>c</u>omplex for <u>p</u>romoting <u>e</u>xit-<u>1</u> (ESCPE-1) complex, which consists of dimers of SNX1 or SNX2 with SNX5 or SNX6, has also emerged as a highly specific proximity interactor of ARL14 (Fig. 7A). SNX1 was identified among the preys with high spectral counts and elevated Log2 fold change compared to the mean of the negative controls in both HEK293 and HeLa cells (Fig. 5D). To corroborate the BioID data, PLA were conducted on FlpIn T-Rex HeLa cells overexpressing expressing wild-type ARL14-GFP in an inducible manner. These experiments demonstrated that endogenous SNX1 (Fig. 7B) and SNX6 (Fig. S7A) are proximal to ARL14, indicating a possible association with the ESCPE-1 complex.

**Figure 7.**
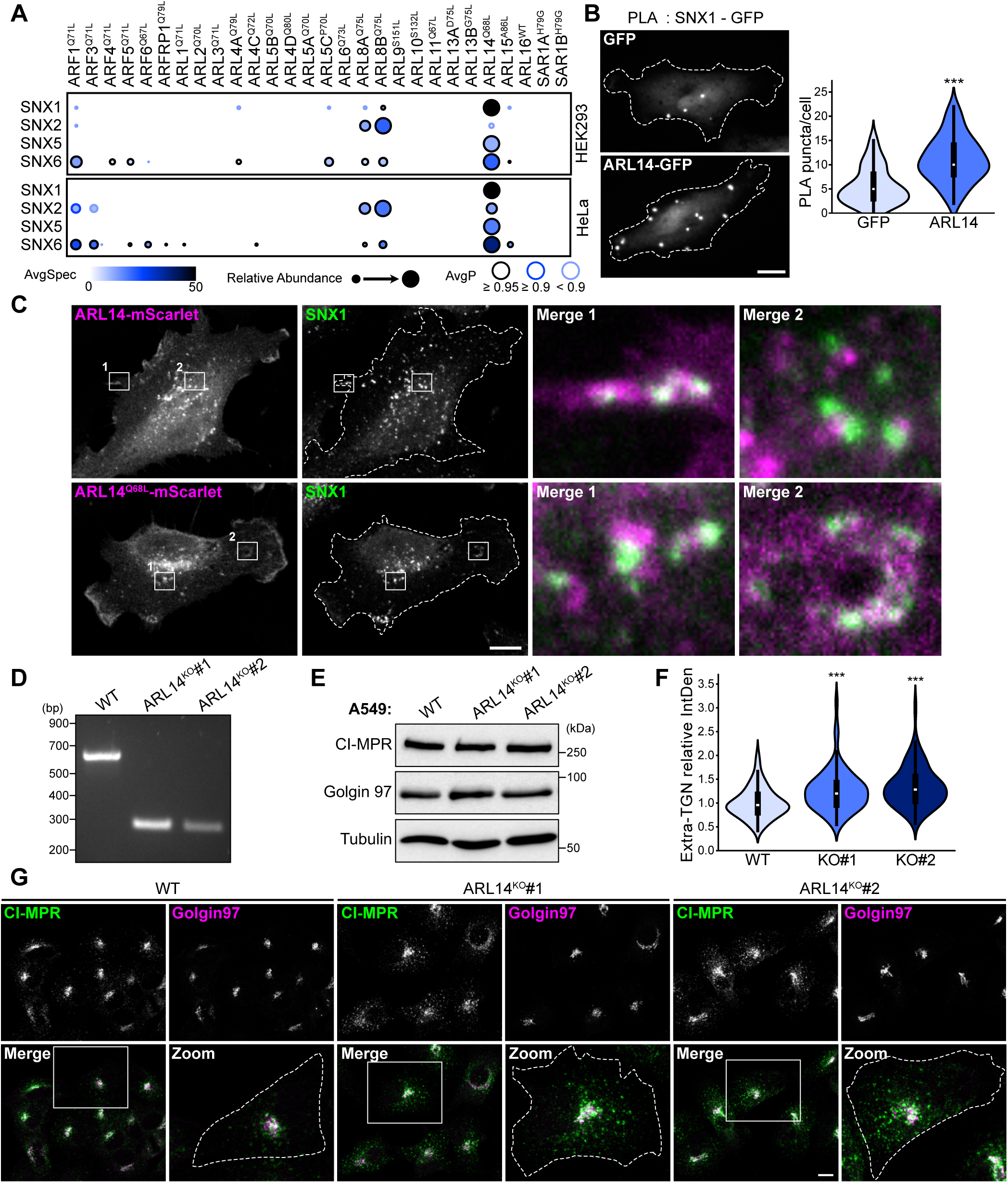
ARL14 is involved in CI-MPR retrograde trafficking through ESCPE-1. **(A)** Dotplots of the BioID data demonstrating the proximity interactions of the ESCPE-1 complex components (SNX1, SNX2, SNX5, SNX6) with ARF GTPases in FlpIn T-REx HEK293 and HeLa cells. Darker circle color represents higher spectral counts, circle size represents relative abundance, and circle outline corresponds to the indicated AvgP. **(B)** Proximity Ligation Assay (PLA) was conducted on endogenous SNX1 and overexpressed wild-type ARL14-GFP in FlpIn T-REx HeLa cells. Cells expressing GFP alone were used as a control. Bar, 10 µm. Violin plot showing the quantification of the number of PLA puncta per cell for each condition. Data are representative of n=3 independent experiments (total of 90 cells per condition). *P* value was calculated by Student *t*-test; \*\*\**P*< 0.0001. **(C)** SNX1 immunostaining in FlpIn T-REx HeLa cells expressing wild-type ARL14-mScarlet or the Q68L constitutively active variant. Merged images represent a ∼8.7x magnification. Bar, 10µm. **(D)** Validation of ARL14 knockout (KO) in A549 cells by PCR. The utilized strategy predicts a band of 610 bp in wild-type (WT) cells and of 280 bp in KO cells. **(E)** Western blot of total proteins extracted from A549 cells, either WT or KO for ARL14, demonstrating no alterations in the expression level of CI-MPR or Golgin97. **(F)** Violin plot showing the quantification of the integrated density (IntDen) outside the TGN relative to the control, for each condition presented in (G). Data are representative of n=3 independent experiments (total of 120 cells per condition). *P* value was calculated by one-way ANOVA, followed by Bonferroni’s test; \*\*\**P* < 0.0001. **(G)** Co-immunostainings was performed using anti-Golgin97, as TGN marker, and anti-CI-MPR in A549 cells either WT or KO for ARL14. Dotted lines delineate cell outlines, which are defined using F-actin staining (not shown). Zoomed in images represent a ∼2.2x magnification. The scale bar represents 10µm.

To delineate the subcellular localization of the ARL14 and ESCPE-1 complex, FlpIn HeLa cells expressing wild-type or the potentially active mutant (Q68L) of ARL14-mScalet were fixed and subjected to staining for SNX1 or SNX6. Given the pivotal role of SNX-BAR proteins in both endosome-to-plasma membrane recycling and endosome-to-TGN retrieval (Simonetti et al., 2023), we predicted that ARL14 and SNX would exhibit colocalization at least in one of these sites. These experiments revealed that both the wild-type and the Q68L variant ARL14-mScarlet were partially soluble in the cytoplasm, with some enrichment in punctate structures reminiscent of endosomes as well as at the plasma membrane (Fig. 7C and Fig. S7B). Puncta positive for either wild-type or Q68L ARL14-mScarlet were frequently appeared adjacent to SNX1 or SNX6 puncta, as depicted in the magnified images (Fig. 7C and Fig. S7B). Furthermore, we observed an accumulation of SNX1 and SNX6 puncta along domains of the plasma membrane enriched for wild-type or Q68L ARL14-mScarlet (Fig. 7C and Fig. S7B). These findings suggest that (i) the Q68L mutation, introduced to constitutively activate ARL14 for the BioID, did not alter ARL14 localization, and (ii) indicate the proximity of ARL14 to the ESCPE-1 complex.

In addition to its roles in phospholipid recognition and membrane tubulation, ESCPE-1 also acts as a recognition and sorting module for a specific subset of transmembrane proteins transiting in the endosomes (Simonetti et al., 2019). One well-studied cargo of ESCPE-1 is the Cation-Independent Mannose-6-Phosphate Receptor (CI-MPR) (Fig. S7C). Depletion of SNX5 and SNX6 was previously demonstrated to reduce CI-MPR retrieval from endosomes to the TGN network, resulting in an accumulation of CI-MPR outside the TGN (Kvainickas et al., 2017, Simonetti et al., 2017). We were able to replicate these findings in HeLa cells using pools of small interfering RNAs (siRNAs) targeting *SNX5* and *SNX6* (Fig. S7D and F). To investigate the potential role of ARL14 in ESCPE-1 mediated trafficking, we utilized CRISPR/Cas9 to generate two *ARL14* knock-out cellular models. We chose to work with HeLa cells, which exhibit low expression of ARL14 and are a direct link to the BioID screens. Additionally, we employed A549 lung cancer cells, previously shown to express ARL14 and in which ARL14 was demonstrated to enhance cell proliferation, migration and invasion (Guo et al., 2019). Following transfection with *ARL14*-specific gRNAs, we isolated clones and confirmed the efficiency of *ARL14* genomic deletion through genomic DNA sequencing (HeLa, Fig. S7G) and PCR amplifications (A549, Fig. 7D). One clone in HeLa and two independent clones in A549 were validated and selected to further investigate the impact of ARL14 on CI-MPR localization. Since the CI-MPR localization assay relies on staining of CI-MPR and Golgin97, we assessed if the absence of ARL14 would lead to variation in the expression levels of CI-MPR or Golgin97, which was not the case (Fig. 7E). Next, by comparing the localization of CI-MPR in or outside of the TGN marked by Golgin97 in control or ARL14 knockout cells, we demonstrated that ARL14 is crucial for the retrieval of CI-MPR to the TGN (A549, Fig. 7F and G; HeLa, Fig. S7H and I). These findings collectively support a previously unknown functional association between ARL14 and the ESCPE-1 in regulating endosomal trafficking.

## DISCUSSION

The ARF and ARL GTPases regulate a broad array of cellular processes, with dynamic protein-protein interactions occurring in membrane-rich compartments being indispensable. Although classical ARFs have been extensively investigated, the localization and functions of ARL proteins remain less defined. In this study, we systematically elucidated the proximity interactomes of ARF and ARL members using BioID, a robust tool for identifying stable or transient protein proximity interactions within their native milieu. Our findings significantly broaden the repertoire of candidate interactors associated with ARF and ARL proteins. These data, available for public access, hold considerable potential for the research community to formulate novel hypotheses. Our investigations represent just the initial step in extracting new insights from this large dataset, particularly regarding the subcellular compartments on which each ARFs and ARLs exerts their functions, their proximity interactors, and new functional implications for the understudied members, such as ARL10 and ARL14.

Our approach has certain limitations, particularly in our requirement to express tagged proteins, which can lead to the creation of artifacts, hinder some interactions, or alter functionality. Additionally, the localization of the GTPases can be skewed to favor one organelle over another. For instance, ARL4D was predominantly localized in the nucleus, despite that its localization to the plasma membrane can be stabilized by binding to the chaperone-like protein HYPK (Lin et al., 2022). It also remains important to validate BioID interactions using additional methods. Moreover, our analysis of ARFs and ARLs was restricted to HEK293 and HeLa cells, possibly missing unique functions and additional interactors specific to specialized cell types. For example, as reported in this study, ARL14 is predominantly express in gastrointestinal cells.

As the endogenous activation mechanisms for most ARF/ARL GTPases are unknown, we performed the BioID screen using mutants that are presumed constitutively active, based on homology to the KRAS^Q61L^ variant. This approach proved highly valuable in enriching our datasets with novel effectors and regulatory proteins, and in helping obtain organelle-specific functions, though undoubtedly ARF/ARL cycling is important for function completion (Klein et al., 2006). Our activating mutations were designed by sequence homology to RAS and well-characterized classical ARFs, but working with unstudied ARLs raises the caveat of uncertainty regarding the effect of these mutations on cycling, likely exacerbated by their weaker affinity for nucleotides compared to RAS and RHO GTPases (Jacobs et al., 1999, Linari et al., 1999, Hanzal-Bayer et al., 2005). Indeed, our HPLC approach provides the first ever glimpse at ARF GTPase nucleotide loading in cells, and the data suggest neither the wild-type proteins or “activated” mutants behave in a manner completely analogous to classical RAS. Previous work suggests some ARF interactions occur only in their GDP-bound state (Donaldson and Jackson, 2011), which supports our finding that several ARF GTPases are not substantially GTP-loaded.

Our NMR and ITC experiments begin to elucidate the biochemical properties of a subfamily of ARFs comprising ARL4A/C/D, ARL11 and ARL14 (Colicelli, 2004). It appears unlikely that ARL14 is regulated by GDP/GTP cycling as it has no affinity for either nucleotide, while ARL11 has an unusually enhanced affinity for GTP. This remains significantly lower than classical RAS proteins that bind both GTP and GDP with low picomolar affinity (Ford et al., 2009). NMR spectra did not present the well-dispersed peaks characteristic of RAS and RHO GTPases, and it is probable that ARL11 and ARL14 are purified from bacteria in an apo state. Previous work suggests this is also true of ARL4A/C (Jacobs et al., 1999), and these results are consistent with our HPLC data. While ARF proteins are difficult to express and purify, future efforts must fully characterize the biochemical activity of all ARF/ARLs to allow improved design of constitutively active and inactive variants.

An objective of this study was to attribute potential functions to less characterized members of the ARF family. Among these, the understudied ARL10 caught our attention due to its localization to mitochondria, as revealed by the BioID data. We determined that this localization is mediated by a distinctive transmembrane domain followed by a polybasic region. Both BioID and complementary AP-MS studies emphasized ARL10’s potential role in mitophagy, mitochondrial fission, and various metabolic processes. Furthermore, our observations also indicate an association between ARL10 and the mTORC2 complex, as evidenced in both ARL10^S321L^ BioID and AP-MS experiments. The preferential interaction of the mTORC2 complex with wild-type ARL10 was confirmed through co-immunoprecipitation with RICTOR. Given ARL10’s localization at the mitochondria, it may provide insights into the function and regulation of the small active pool of mTORC2 previously observed at this location (Ebner et al., 2017). Furthermore, our ARL10 BioID data also identified the presence of several peroxisome-resident proteins. This was corroborated by co-immunostaining, confirming the existence of a pool of ARL10 at peroxisomes. Peroxisome and mitochondria are closely interconnected organelles that communicate through small vesicles (Neuspiel et al., 2008). Because of the localization of ARL10 to mitochondria and peroxisomes, and its proximity to proteins involved in mitochondrial dynamics, an attractive hypothesis for future research is that ARL10 may control vesicular traffic between these organelles.

While our study was nearing completion, Li *et al*. published an interactome of the ARF family, opting to use a miniTurboID approach, biotin labeling for only 15 minutes, targeting 25 constitutively active members of the family (excluding ARL4A, ARL5C, ARL9 and ARL13A). In contrast to our method of expressing baits in a tetracycline-inducible manner, they employed stable overexpression by infecting HEK293A cells. Based on our experience with RHO proteins, we chose to perform BioID with baits expressed in an inducible manner in HEK293 and HeLa cells, to mitigate toxicity or gene expression alterations resulting from stable and uncontrolled overexpression of constitutively active GTPases. The Li *et al*. study and our work presented here are complementary and demonstrate that short versus long-term labeling can yields unique insights. Short labeling highlights frequently occurring interactions, while longer labeling captures more transient interactions, increases the overall number of proximal interactors, and reveals interactions dependent on specific cell states (*e.g.* during specific stages of the cell cycle). We compared the hits from Li *et al*. to our datasets obtained in HEK293 cells for the 24 common baits. We estimate approximately 350 shared interactions, leaving nearly 1400 new proximity interactors identified by our BioID. A limitation of the Li *et al*. approach is the lower spectral counts observed (due to short labeling time), resulting in very low correlation coefficients among replicates (some as low as 0.4). In our study, all replicates show correlation coefficients above 0.9. Also, the Li et al. study does not provide statistical data to support the confidence of the reported proximity interactions. In contrast, we rigorously utilized non-specific baits (BirA*, BirA*-eGFP and BirA*-eGFP-CAAX) to filter noise from our ARF/ARL baits, and we provide statistical confidence for all our data. In conclusion we believe that these two studies together represent a significant advancement in obtaining an accurate representation of the entire ARF family interactome.

Like us, Li *et al*. noted the previously unexplored interaction between PLD1 and ARL14. While they demonstrated that ARL14 is a PLD1 activator *in vitro*, we employed an *in cellulo* approach, providing further evidence for ARL14-mediated activation of PLD1, with the two approaches thereby strengthening the conclusion that PLD1 is a *bona fide* effector of ARL14. While Li *et al*. defined the essential region in PLD1 for ARL14-binding, we conducted functional studies that led to the identification of the ESCPE-1 complex as a potential effector of ARL14. Notably, our experiments demonstrated a reduction in CI-MPR retrieval in the Golgi of ARL14-null A549 and HeLa cells. We identified SNX1, SNX2, SNX5 and SNX6 as relatively specific proximal interactors of ARL14, but only SNX1 was identified in the ARL14 miniTurboID of Li *et al*.. It is worth mentioning that SNX1, SNX2, SNX5 and SNX6 are also associated with the SNX-BAR Retromer complex, which includes the heterotrimer VPS35– VPS26–VPS29 and a dimer of SNX1 or 2 with SNX5 or 6 (Bonifacino and Hurley, 2008). Although the VPS trimer was not identified in either the ARL14 BioID or miniTurbo experiments, it cannot be excluded that ARL14 connect with this complex; further experimentations will be required to investigate this. Based on our findings, we propose the postulate that the ESCPE-1 complex and PLD1 might operate in collaboration to generate endosomes. ARL14 could insert into endosomal membranes, inducing local positive curvature, thereby attracting ESCPE-1 as coats, leading to tubulation and concentration of cargos. Concurrently, ARL14 could recruit and activate PLD1 to produce PA, enhancing membrane curvature for endosome formation. Additionally, our study reveals the restricted expression of ARL14 in gastrointestinal cells of mice. Fully characterizing the role of ARL14, and its interactors PLD1 and ESCPE-1, in specialized cells of the intestines and within the context of lung cancer, where these protein-protein interactions may play crucial roles in secretory functions, represents a biologically relevant future area of investigations.

Overall, our study underscores the potential of our dataset in uncovering new functions and regulatory mechanisms of ARF and ARL proteins. Our BioID data is accessible on our website, serving as a valuable resource for the research community interested in exploring interactors of ARF and ARL proteins.

## METHODS

### Antibodies, plasmids, siRNAs, primers, cDNA and cloning

The antibodies used in this study are detailed in Table S2, while plasmids are catalogued in Table S3, siRNAs in Table S4 and primers in Table S5. The NCBI Reference Sequence numbers of the complementary DNA (cDNA) of ARF/ARL genes used in this study are listed in Table S1. All coding sequences were synthesized as previously described (Patel et al., 2011), with the exception of ARL6, ARL9 and ARL16, for which sequences were obtained from NCBI and synthetized with addition of Gateway compatible AttB sites (Twist Bioscience). The Gateway cloning system (Invitrogen) was used to recombine the cDNA sequences into the entry vector (pDONR221). Subsequently, each ARF/ARL cDNA was recombined into the expression vector pcDNA5-pDEST-FRT-BirA*-Flag-CT (Couzens et al., 2013). For certain proteins, alternative destination vectors were used to clone in frame fluorescent protein coding sequences, namely pcDNA5-pDEST-FRT-eGFP-CT (Kean et al., 2011) was used for ARL10 and ARL14 and pcDNA5-pDEST-FRT-mScarlet-CT for ARL14. To construct pcDNA5-pDEST-FRT-mScarlet-CT, *mScarlet* coding sequence was amplified by PCR, with pMaCTag-Z11 (Addgene# 120054) as a template DNA, using primers engineered with KpnI/XhoI cloning sites, such that the mScarlet cDNA replaced the eGFP sequence in pcDNA5-pDEST-FRT-eGFP-CT after digestion with KpnI and XhoI and ligation.

### Cell culture and transfections

Flp-In^TM^ T-Rex^TM^ HEK293 (Thermo Fisher), Flp-In^TM^ T-Rex^TM^ HeLa (gift from S. Taylor, University of Manchester UK), HEK293T (ATCC), and HeLa (ATCC) cell lines were maintained at 37°C and 5% CO_2_ and grown in Dulbecco’s modified Eagle’s medium (DMEM) supplemented with 10% fetal bovine serum (FBS) (F1051, Sigma-Aldrich) and 1% penicillin– streptomycin (P/S) (450-201-EL, Wisent). Stable cell lines were generated as previously described (Nahle et al., 2022) and were grown in DMEM supplemented with 10% tetracycline-free FBS (081-150, Wisent) and 1% P/S. To induce protein expression, stable cell lines were treated with 1 ug/mL of tetracycline with or without 50 µM biotin supplementation. siRNA transfections were performed using Dharmafect-1 (GE Healthcare) and 80nM of indicated siRNA. Plasmid DNA transfections were performed with Lipofectamine^TM^ 3000, according to the manufacturer (L3000001, Thermo Fisher).

### Protein expression purification

GST-tagged ARL11 (1-196) and ARL14 (1-192) were expressed and purified from *Escherichia coli* (BL21-DE3-codon+) cells cultured in LB media. For NMR, isotopically-labeled proteins were obtained by growing BL21-DE3-codon+ cells in M9 minimal media [50 mM Na_2_HPO4, 22 mM KH_2_PO4, 8.5 mM NaCl, 1% glucose, 1 mM MgSO_4_, 0.1 mM CaCl2, 38 µM thiamine, pH 7.4], and for ^15^N-uniformly labeled protein samples, 1 g of [^15^N]NH_4_Cl was added. Cells were grown at 37 °C until reaching an optical density of 0.6-0.8, followed by induction with 250 µM of isopropyl-β-D-thiogalactopyranoside (IPTG) (800-050-EG, Wisent). After induction, the temperature was decreased to 18 °C and cells grown overnight. Harvested cells were lysed in a buffer containing 20 mM Tris-HCl pH 7.5, 150 mM NaCl, 10% glycerol, 0.4% NP-40, protease inhibitors (Roche) and 2 mM dithiothreitol (DTT), followed by sonication. Each lysate was clarified by centrifugation and next incubated with glutathione sepharose resin (17-0756-04, Cytiva) at 4°C for 1-2 hours. Subsequently, bound proteins were cleaved from the resin, without elution, using thrombin protease (91-035-100, BioPharm). The recovered proteins after cleavage underwent size exclusion chromatography using a Superdex S75/300 column (GE Healthcare) in phosphate-based buffer [200 mM sodium phosphate pH 7.4, 15 mM HEPES pH 7.4, 10 mM MgCl_2_, 5 mM KCl, 2 mM DTT].

### HPLC Nucleotide Analysis

One 15 cm plate of HEK293T cells was transfected with the indicated plasmids using Opti-MEM^TM^ (31985062, ThermoFischer) and Polyethylenimine (23966-1, Polysciences), followed by harvest after 48 hours. To isolate GFP-tagged GTPases, the cells were suspended in lysate buffer [20 mM Tris-HCl pH 7.5, 150 mM NaCl, 5 mM MgCl_2_, 10% glycerol, 1% NP-40, 1% Triton-100X, 1 mM DTT, protease inhibitors], then incubated on ice for 15 min. Each lysate was then cleared at 15000 rpm for 10 min at 4°C, and the cleared lysate was incubated with NHS beads (GE17-0906-01, Cityva) pre-conjugated with anti-GFP nanobodies for 30 min. Beads underwent a single buffer wash step to minimize the loss of weakly bound nucleotides using the following buffer [20 mM Tris-HCl pH 7.5, 150 mM NaCl, 5 mM MgCl_2_,1 mM DTT]. Following this, the beads were resuspended in 200 µL of wash buffer and boiled at 95°C for 6 min, followed by centrifugation at 15000 rpm for 10 min. The supernatant, containing released nucleotides, was filtered through pre-washed PVDF centrifugal filters (0.22 µm PVDF centrifugal filter (UFC30GV25, Millipore)). The flow through was used for HPLC analysis, conducted using an Agilent 1100 Series HPLC and C18 column (Eclipse XDB-C18 5 μm, 4.6×150 mm (Agilent)). The running phase was prepared in 500mL: 6.29 g KH_2_PO_4_, 1.49 g tetrabutylammonium bromide, and 6.8% acetonitrile in water. Each sample was run in the mobile phase at a flow rate of 0.95 ml/min for 9 min at 25°C, with absorbance detected at 252 and 254 nm.

### BioID and mass spectrometry injections

BioID experiments were performed as previously described in (Bagci et al., 2020), with additional protocols for BioID and MS injection further detailed in (Nahle et al., 2022). Briefly, cells were lysed using RIPA lysis buffer [1% NP-40, 50mM Tris-HCl pH 7.4, 0.1% SDS, 150mM NaCl, 0.5% sodium deoxycholate, 1mM EDTA pH 8], followed by sonication. Biotinylated proteins were isolated from cleared lysates by incubating with 70 µl of streptavidin beads (17-5113-01, GE Healthcare) at 4°C for 3h. Following successive washes with lysis buffer and ammonium bicarbonate (AB0032, Biobasic), on-beads digestion was performed overnight with 1 ng/µL of trypsin (T6567, Sigma-Aldrich), followed by an additional 2h incubation with 1 ng/µL of fresh trypsin. The digested peptides were recovered from the mix containing beads by washes with water. Tryptic digestion was terminated by adding formic acid (94318, Sigma-Aldrich) to a final concentration of 5% to end. Peptides were then dried down in a SpeedVac, resuspended in 15 µl of 5% formic acid, and stored at −80°C. Samples generated from HEK293 cells were analyzed with an LTQ-Orbitrap Velos (Thermo Fisher) while samples from HeLa cells with a Q Exactive (Thermo Fisher). Peptide samples were loaded into a 75 μm i.d. × 150 mm Self-Pack C18 column installed on the Easy-Easy-nLC 1000 system (Proxeon Biosystems). Elution of peptides was performed with a two-slope gradient at a flow rate of 250 nL/min, from 2% to 35% in 100 min and then from 35 to 80% in 10 min transferring from buffer A [0.2% formic acid] to buffer B [90% acetonitrile/0.2% formic acid]. Ionisation of peptides was achieved using a Nanospray Flex Ion Source on both instruments. Acquisition in the Orbitrap was made with a full-scan MS survey scanning (m/z 360–2,000) in profile mode with a resolution of 60,000, an AGC target at 1E6 and a maximum ion time of 100 ms for both machines. Nanospray and S-lens voltages were set to 1.3–1.8 kV and 50 V, respectively. Capillary temperature was set to 250°C. On the LTQ Orbitrap Velos, the 11 most intense peptide ions were fragmented by collision-induced dissociation fragmentation and MS/MS spectra were acquired in the linear ion trap with an AGC target at 1E4 and a maximum ion time of 100 ms. MS/MS parameters were: normalized collision energy, 35 V; activation q, 0.25; activation time, 10 ms and target ions already selected for MS/MS were dynamically excluded for 30s. For the Q Exactive, the 16 most intense peptide ions were fragmented by higher-energy collisional fragmentation and MS/MS spectra were acquired in the Orbitrap with a resolution of 17,500, an AGC target at 1E5 and a maximum ion time of 50 ms. MS/MS parameters were: normalized collision energy at 27 and target ions already selected for MS/MS were dynamically excluded for 7s.

### Mass spectrometry data analyses

The raw mass spectrometry files underwent analysis using the X! Tandem (Craig and Beavis, 2004) and Mascot search engines, through the iProphet pipeline (Shteynberg et al., 2011) integrated in the Prohits analysis suite (Liu et al., 2010). The searches were performed using the Human RefSeq protein sequence database (2017 version) supplemented with “common contaminants” from the Max Planck Institute (http://maxquant.org/downloads.htm), the Global Proteome Machine (GPM; http://www.thegpm.org/crap/index.html), and along with decoy sequences. Mascot parameters were configured with strict trypsin specificity (allowing for two missed cleavage sites) and Oxidation (M) and Deamidation (NQ) considered as variable modifications. The mass tolerances for precursor and fragment ions were set to 10 ppm and 0.6 Da, respectively, with peptide charges of +2, +3 and +4 being considered. The resulting X! Tandem and Mascot search results were individually processed by PeptideProphet (Keller et al., 2002), and peptides were assembled into proteins using parsimony rules first described in ProteinProphet (Nesvizhskii et al., 2003) using the Trans-Proteomic Pipeline (TPP). TPP settings were as follows: -p 0.05 -x20 -PPM –d“DECOY”, iprophet options: pPRIME and PeptideProphet: pP.

Samples reproducibility among biological replicates was assessed in R (r-project.org) by performing multidimensional scaling with the limma package (Ritchie et al., 2015) and Spearman correlations, with the stats package using spectral counts of identified preys. We discarded and reacquired biological replicates displaying Spearman correlations lower than 0.9 and/or clustered poorly among biological replicates on a MDS plot.

### Interactions scoring

To calculate statistics on the interactions, we used SAINTexpress (Teo et al., 2014) (version 3.6.1) on proteins with iProphet protein probability ≥ 0.9 and unique peptides ≥ 2. Proteomics datasets (ARFs BioID screens in HEK293 and HeLa cells) were compared separately against their respective negative controls. These controls comprised of pulldowns performed on both cell lines expressing either the eGFP-BirA*-Flag vector or the empty vector (BirA*-Flag alone), each in biological duplicates. SAINT analyses were performed without control or bait compression. To infer ARFs’ high confidence interacting proteins, we applied, in addition to the SAINT average probability (AvgP) ≥ 0.95 threshold (more stringent than a BFDR ≤ 0.01 threshold), a combination of two additional filters: the preys’ Average Spectral count (AvgSpec) and the preys’ CAAX ratio (Preys AvgSpec/CAAX AvgSpec). For each the cell line, the cutpoints of both additional filters were estimated by cumulative distribution (CDA) and ROC sensitivity analyses. After identifying ARFs and ARLs physical recall interactions, as reported by the human BioGrid database (v 4.4.210) (Chatr-Aryamontri et al., 2017), we performed CDA and ROC sensitivity analysis in R with the mltools and cutpointr packages, respectively. To gain precision, we limited these analyses to ARF1, ARF6, SAR1A and SAR1B, which have been, up to this date, the most characterized proteins of the ARF family. From the results of these approaches, we reasoned that AvgSpec thresholds of 4.5 and 6, with a CAAX ratio of 1.7, were the most appropriate for the HEK293 and HeLa datasets, respectively. Unfiltered contaminants, such as Keratin, BirA*, carboxylases and ß-galactosidase were manually removed. To assess the specificity of each interaction among the ARFs interactome, we calculated for each cell line, their WD-score by using the SMAD R package (Sowa et al., 2009). Next, we pooled the interactions specificity of both cell lines by summing their respective WD-scores, giving us a “WD-Summed score” that we referred to as WDS score. The benefit of this metric is to increase our capacity to identify ARF/ARL interactors of high confidence, either from interactions shared by both cell lines or from unique cell line interactions. We then exploited this WDS score metric by building a heatmap of ARFs and ARLs effectors with the R complexheatmap package (Gu et al., 2016). To do so, we selected potential ARFs and ARLs effectors by removing GAPs and GEFs from filtered ARF/ARL interactions. Then we calculated Canberra distances between filtered preys and Pearson correlations between baits, followed by using the Ward.D method to extract 16 clusters of preys and 13 clusters of baits. The optimal number of clusters was determined using the silhouette method. Similarly, heatmaps of GAPs and GEFs were also created without cluster extraction. For each of the clusters of ARFs and ARLs effectors, we performed an over-representation analysis of GO BP in R using the gprofiler2 package (Kolberg et al., 2020). The complete lists of interactions with scores and annotations are available online at the following website: http://prohits-web.lunenfeld.ca.

### Bioinformatic and network analyses

A proportional area chart was created using the ggplot2 package in R. Analysis of each ARF/ARL subcellular localization was executed separately on the Cell Map website (cell-map.org), with the results filtered to the top three subcellular localizations with the shortest distances. Graphical representations of protein-protein networks were generated using Cytoscape (Shannon et al., 2003) (version 3.9.1). The illustration of the ARFs/ARLs subcellular localizations was inspired by the animal cell drawing from the SwissBioPics website (swissbiopics.org). Dotplot representations of interactions were generated at the ProHits-viz website (prohits-viz.org). Enrichment maps of Gene Ontology “Biological Processes” (GO BP) were generated in Cytoscape with the EnrichmentMap app (v 3.3.4) (Merico et al., 2010), by loading overrepresented GO BP estimated with fdr correction by gProfiler (Raudvere et al., 2019). CBNP normalized values were calculated as previously described (Cozzolino et al., 2020).

### Immunofluorescence staining, imaging, and analyses

Cells were fixed with 3.7% formaldehyde in CSK buffer [100 mM NaCl, 300 mM sucrose, 3 mM MgCl_2_, 10 mM PIPES, pH 6.8] for 15 min, followed by permeabilization with 0.2% Triton X-100 in PBS for 5 min. Subsequently, cells were incubated with primary antibody in the wash buffer [1% BSA, 0.1% Triton-X100 in TBS], washed with wash buffer, and incubated, for 30 min, with the appropriate secondary antibody. Where indicated, Hoechst, phalloidin and/or streptavidin were simultaneously incubated with secondary antibodies. Afterward, cells were washed with the wash buffer and then with water. Coverslips were mounted with Mowiol. Images were acquired with a Carl Zeiss LSM700 laser scanning confocal microscope (Carl Zeiss MicroImaging, Jena, Germany) equipped with a plan-apochromat ×63/1.4 numerical aperture objective and operated with ZenBlack 2009. Fiji (Schindelin et al., 2012) (v2.0.0-rc-69/1.52k) was used for analyses. The fluorescence intensity of CI-MPR was quantified by measuring CI-MPR integrated density across the whole cell, excluding the TGN delineated by the Golgin97 staining.

### Co-immunoprecipitation and affinity purification-mass spectrometry

To enrich for ARL10-associated protein proteins, FlpIn HeLa ARL10-GFP cells were lysed in digitonin buffer [50 mM HEPES pH 7.4, 120 mM NaCl, 0.5% digitonin, 0.25 mM NaVO_3_, 1 mM PMSF, 10mM NaF] supplemented with EDTA-free Protease Inhibitor Cocktail (4693132001, Sigma-Aldrich). 1mg of cleared cells lysate was incubated with 10 µL of GFP-selector affinity resin developed by NanoTag Biotechnologies (N0310, Synaptic Systems) for 2h at 4°C. Beads were washed three times with lysis buffer, resuspended in 30 µl of Laemmli buffer [5% SDS, 0.1 mM Tris pH 6.8, 140 mM β-mercaptoethanol, 25% glycerol] and denatured for 5 min at 95°C for western blotting.

For affinity purification-mass spectrometry of the ARL10-GFP associated proteins, the protocol above was followed with the exception that after the three washes of the pulled down GFP-selector beads with lysis buffer, two additional washes with 50 mM HEPES/120 mM NaCl were performed to remove detergents. Samples were then reduced with 9 mM dithiothreitol at 37°C for 30 minutes and, after cooling for 5 minutes, alkylated with 17 mM iodoacetamide at room temperature for 30 minutes in the dark. Prior to protein digestion, a Protein Aggregation Capture using Hydroxyl beads (ReSyn Biosciences) was performed to remove SDS and glycerol. The proteins were digested on-beads with Trypsin (Promega) with ratio enzyme:proteins 1:20 in 50 mM ammonium bicarbonate and performed overnight at 37°C. For each sample, the supernatant was transferred into a new sample tube and dried with a vacuum centrifuge (Eppendorf). Samples were reconstituted under agitation for 15 min in 11 µL of 2%ACN-1%FA and loaded into a 75 μm i.d. × 150 mm Self-Pack C18 column installed in the Easy-nLC II system (Proxeon Biosystems). Peptides were eluted with a two-slope gradient at a flow rate of 250 nL/min. Solvent B first increased from 2 to 35% in 120 min and then from 35 to 85% B in 20 min. The HPLC system was coupled to Orbitrap Fusion mass spectrometer (Thermo Scientific) through a Nanospray Flex Ion Source. Nanospray and S-lens voltages were set to 1.3-1.7 kV and 50 V, respectively. Capillary temperature was set to 225°C. Full scan MS survey spectra (m/z 360-1560) in profile mode were acquired in the Orbitrap with a resolution of 120,000 with a target value at 1e6. The 25 most intense peptide ions were fragmented in the HCD collision cell and analyzed in the linear ion trap with a target value at 1e4 and a normalized collision energy at 29 V. Target ions selected for fragmentation were dynamically excluded for 20 sec after two MS2 events.

### NMR sample preparation and data acquisition

Uniformly-labeled [^15^N]ARL11 or [^15^N]ARL14 were diluted to 100-200 µM in phosphate buffer on ice. GDP or GTPγS nucleotides were added in 10-fold molar excess to proteins. The final sample solution contained 100-200 µM of protein, 10-fold excess of nucleotide and 10% D_2_O. Samples were centrifugated at 4°C to remove possible precipitation and transferred to a 3 mm NMR sample tube. One dimension (1D) ^1^H (ZGESGP) and two-dimension (2D) ^1^H-^15^N BEST-HSQC (B_HSQCETF3GPSI) spectra were recorded on an 600 MHz Bruker Avance III magnet equipped with a tuneable ^1^H-^19^F QCIP 5 mm cryoprobe at 298 K after short (∼10 minutes) or long (∼17 hours) incubation periods with nucleotide. Data was processed using NMRFx Analyst and visualized with NMRViewJ.

### Isothermal Titration Calorimetry

The affinity of ARL11 or ARL14 to nucleotides was measured using a MicroCal iTC200 (Malvern). Nucleotide stock solutions were diluted using filtered and degassed phosphate-based buffer (as used for size exclusion chromatography of ARL11 and ARL14). Experiments were carried out at 20°C. Data was processed and visualized using Origin 7 (MicroCal).

### Animal experiments

Mice (*Mus musculus*) were housed in a specific pathogen-free (SPF) facility, and all experiments were approved by the Animal Care Committee of the Montreal Clinical Research Institute (protocol 2021-04 MK), in compliance with the Canadian Council of Animal Care guidelines. ARL14^3xFlag^ knock-in alleles were generated by CRISPR/Cas9 mediated gene editing via the *i*-GONAD procedure (Gurumurthy et al., 2019). Briefly, a Cas9 ribonucleoproteins (RNPs) mix, containing 6 μM Cas9 proteins (S.p. HiFi Cas9 Nuclease V3, 1081061, Integrated DNA Technologies), 30 μM gRNA, and the 30 μM ssDNA (coding for 3x Flag), were delivered by microcapillary injection into ampulla of 6 to 10 weeks old CD-1 pregnant females at 0.7 days post conception, followed by electroporation of the oviduct. Mice that were born were genotyped by PCR, followed by Sanger sequencing, to verify correct targeting. To select gRNA, we used the IDT tool: (https://www.idtdna.com/site/order/designtool/index/CRISPR_CUSTOM). All sequences used are described in Table S5.

For organs collection, male mice of 65 days of age were sacrificed. Organs were dissected, flash frozen and kept at −80°C. Each indicated organ was grinded in liquid nitrogen, with an additional shredding step for muscles, and proteins were then extracted with RIPA lysis buffer supplemented with inhibitors [1% NP-40, 50mM Tris-HCl pH 7.4, 0.1% SDS, 150mM NaCl, 0.5% sodium deoxycholate, 1mM EDTA pH 8, 10 mM sodium fluoride, 1 mM sodium orthovanadate and Protease Inhibitor Cocktail]. Proteins were then quantified from cleared lysates using DC Protein Assay (Bio-Rad), and equal amount of each sample was analyzed by western blot. The top half portion of the blot was used for Coomassie Brilliant Blue R-250 staining, to assess total protein loading, and the bottom half was used for immunoblotting with the anti-Flag antibody to assess ARL14^3xFlag^ expression.

### Proximity ligation assay (PLA)

Cells were fixed and permeabilized as described for immunofluorescence staining. PLA was performed using the Duolink® In Situ Detection Reagents Red (DUO92008, Sigma-Aldrich) using the anti-mouse MINUS (DUO92004, Sigma-Aldrich) and the anti-rabbit PLUS (DUO92002, Sigma-Aldrich) probes. Images were acquired with a DM6 upright microscope (Leica Microsystems Inc., Ontario, Canada) equipped with an ORCAflash 4.0 V.2 monochromatic camera (Hamamatsu Photonics, Bridgewater, NJ, USA). PLA dots were counted using Fiji (Schindelin et al., 2012) (v2.0.0-rc-69/1.52k).

### Measurement of PLD activity using IMPACT

For IMPACT, cells were plated onto a 24-well plate and protein expression was induced overnight with 1 µg/mL of tetracycline. Cells were pre-treated with FIPI (750 nM, 5282455MG, MilliporeSigma) or DMSO control for 30 min, followed by 3-azidopropanol (1 mM) (Yuan et al., 2012) and PMA (100nM, P8139, MilliporeSigma) for 30 min. Cells were washed 3 times with PBS and incubated with BCN-BODIPY (1 µM) (Alamudi et al., 2016) diluted in Tyrode’s-HEPES buffer [135 mM NaCl, 5 mM KCl, 1.8 mM CaCl_2_, 1 mM MgCl_2_, 1mg/mL glucose, 1 mg/mL BSA, 20 mM HEPES, pH 7.4] for 30 min at 37°C. Cells were rinsed twice with PBS and split into 3 wells of a 96-well v-bottom plate. The plate was centrifuged at 1000g for 2 min and cells were rinsed twice with PBS. Cells were fixed with 4% PFA in PBS followed by two additional PBS washes. Cells were resuspended in fluorescence-activated cell sorting (FACS) buffer [0.5% FBS in PBS] for FACS analysis, which was performed on a Thermo Fisher Attune NxT analyzer.

### Statistical analyses

All quantification graphs were derived from Prism (GraphPad Software, v7.0a). Statistical significance was determined using Student t-tests or one-way analysis of variance (ANOVA), followed by Bonferroni’s multiple comparison test where *, ** and *** represented P < 0.05, P < 0.001 and P < 0.0001 respectively. Sample sizes and number of replicates are mentioned in figure legends.

## ACKNOWLEDGEMENT

The authors thank S. Taylor (University of Manchester, UK) for the gift of the Flp-In^TM^ T-Rex^TM^ HeLa cells. We also thank M.A. Frohman for the pCGN-HA-PLD1 plasmid. We recognize the technical assistance and expertise provided by the IRCM core facilities, including the proteomic facility, the microscopy facility (Dr. D. Filion) and the animal facility (S. Riverin).

## DATA AVAILABILITY

The raw BioID proteomic data were deposited on MASSIVE for further exploration: https://doi.org/doi:10.25345/C50K26G6S. The complete lists of preys identified with their respective scores and annotations are also available online at the following website: http://prohits-web.lunenfeld.ca.

---

**Figure S1.**
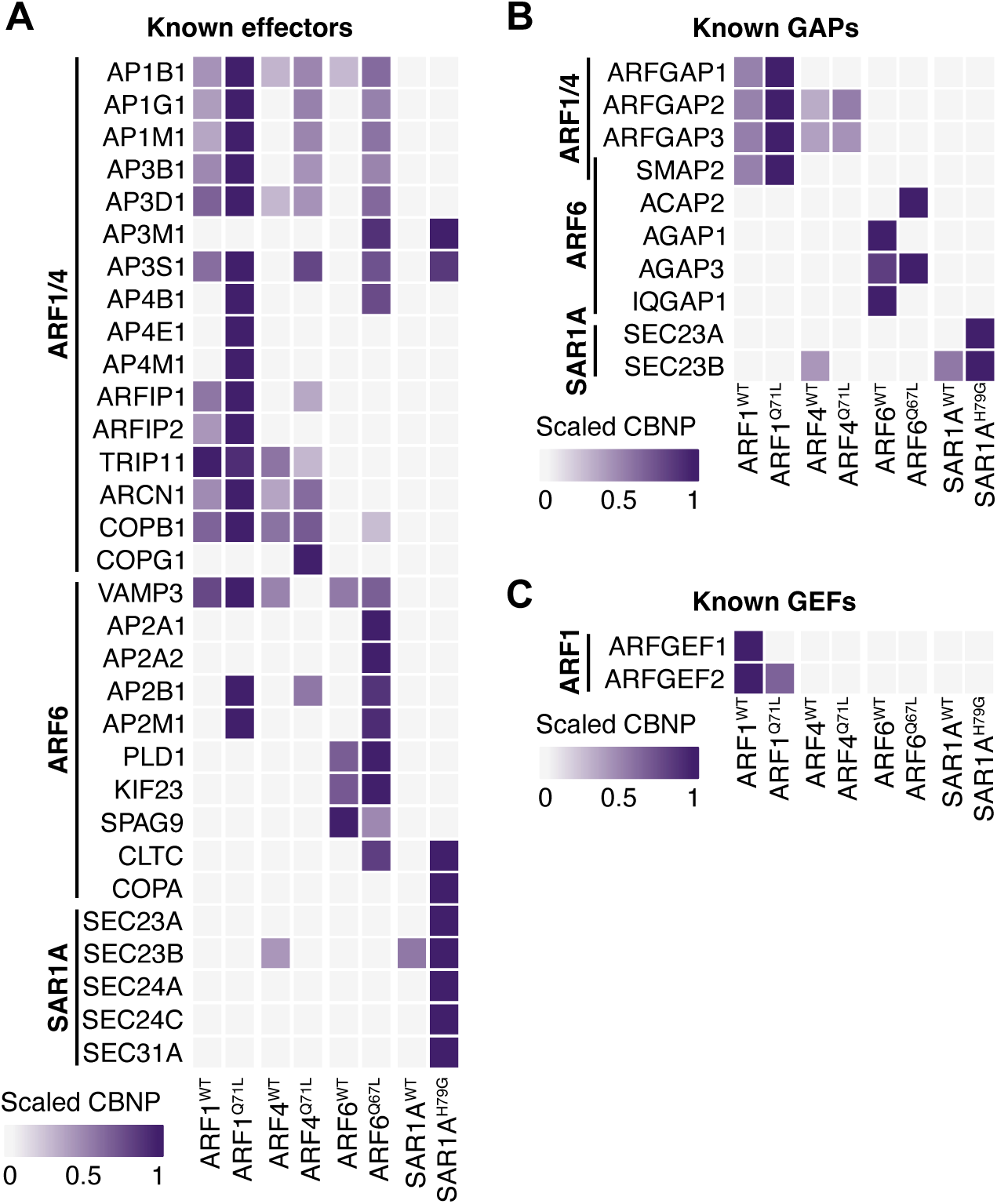
The activated ARF family members allow for effectors and GAPs enrichment. Heatmaps representing the known **(A)** effectors, **(B)** GAPs and **(C)** GEFs identified in BioID performed in FlpIn T-REx HeLa cells expressing the indicated baits. The scale shows the normalized spectral counts (CBNP) where the darker color is indicative of a higher number of spectral counts.

**Figure S2.**
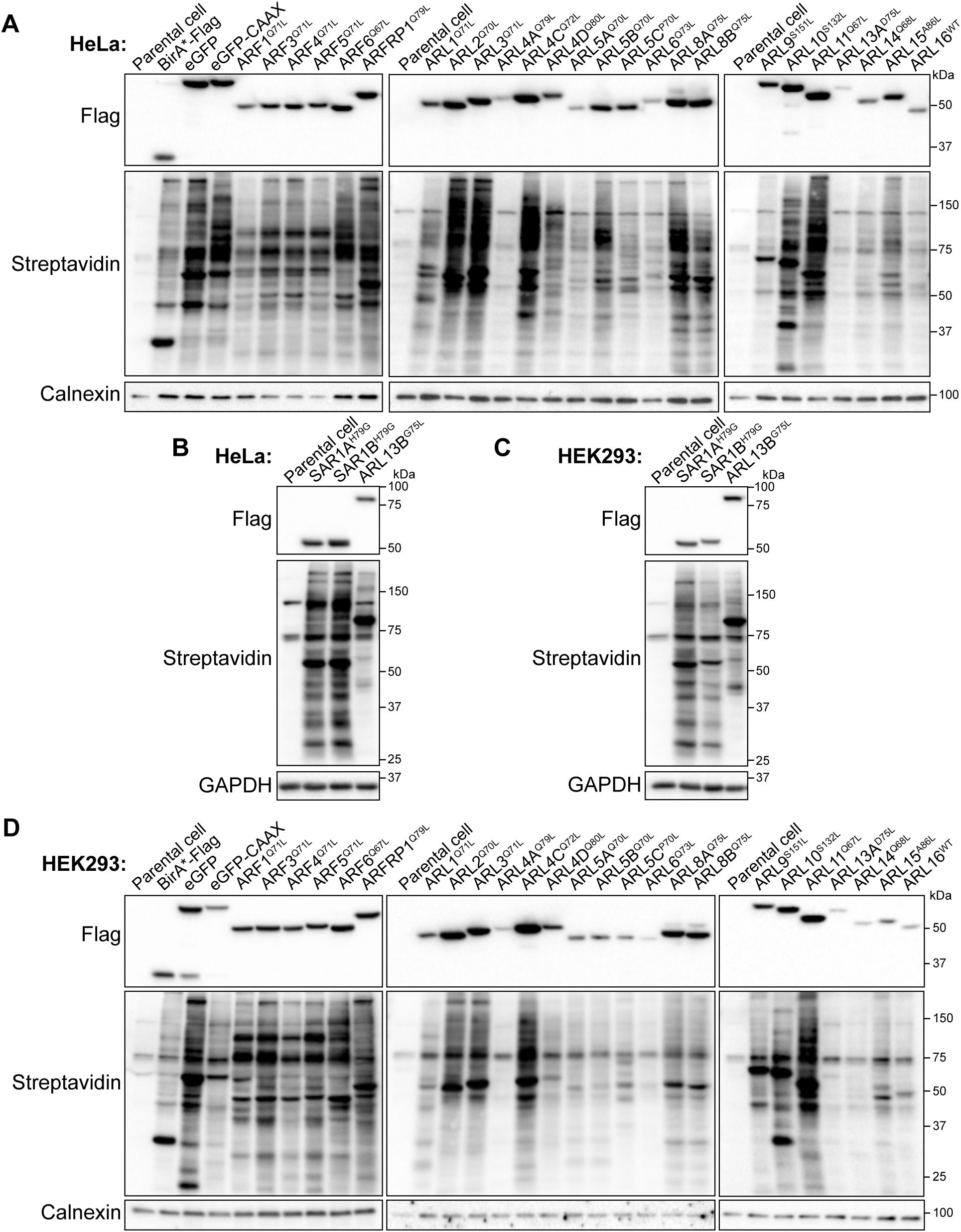
Validation of the expression and biotinylation of ARF-BirA*-Flag cell lines. **(A)** and **(B)** Western blot of FlpIn T-REx HeLa cells expressing the indicated constitutively active ARF-BirA*-Flag construct treated for 24 hours with tetracycline and biotin. Loading controls are calnexin in (A) and GAPDH in (B). Data are representative of n=2 experiments. **(C)** and **(D)** Western blot of FlpIn T-REx HEK293 cells expressing the indicated constitutively active ARF-BirA*-Flag construct treated for 24 hours with tetracycline and biotin. Loading controls are GAPDH in (C) and calnexin in (D). Data are representative of n=2 experiments.

**Figure S3.**
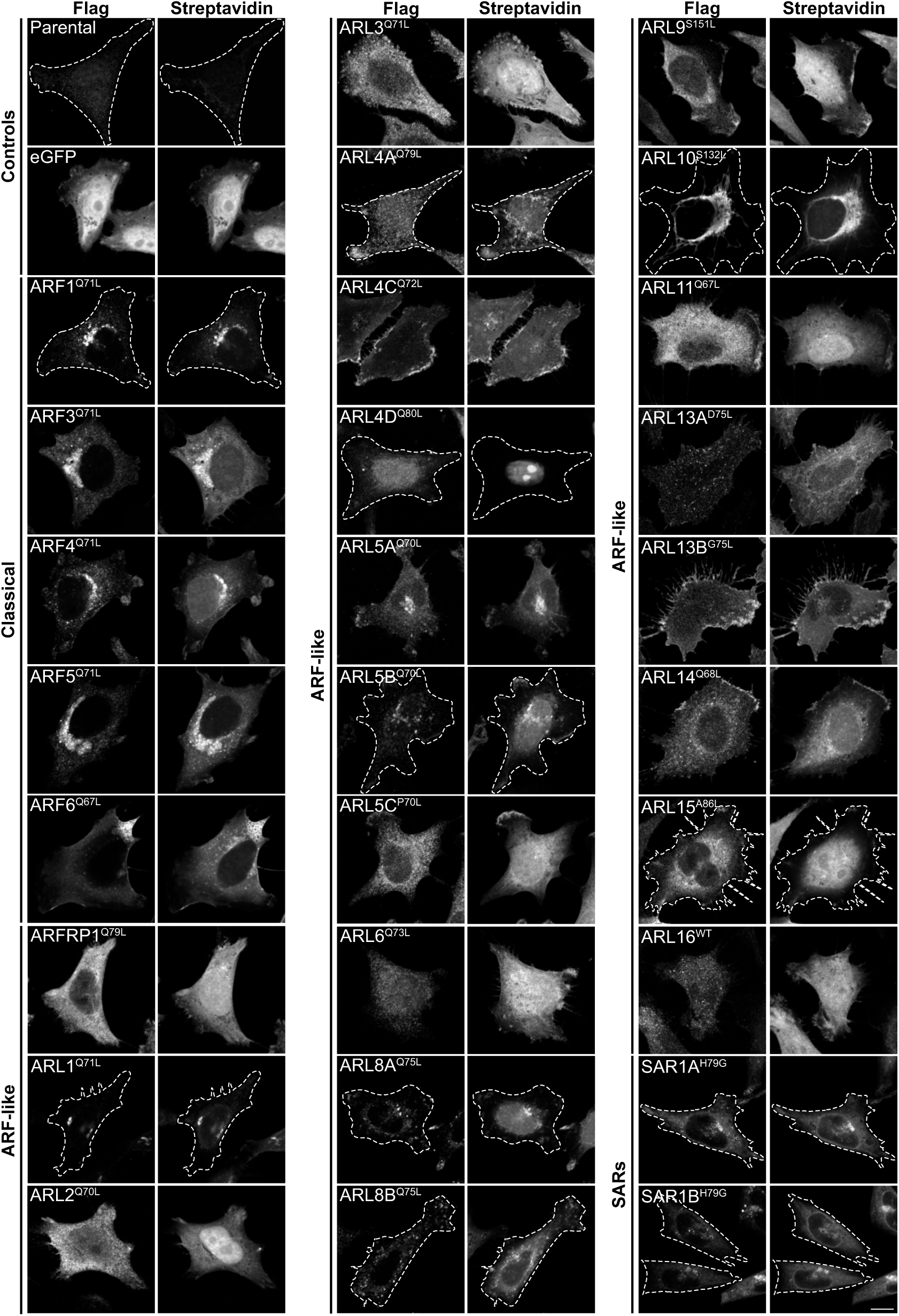
Validation of the expression and biotinylation of ARF-BirA*-Flag HeLa cells. Confocal microscopy images of immunostaining against Flag (Alexa Fluor 488) and streptavidin (Alexa Fluor 647-Streptavidin) in FlpIn T-REx HeLa cells expressing the indicated constitutively active ARF-BirA*-Flag construct or control treated for 24 hours with tetracycline and biotin. Parental cells were used as control. When necessary, F-actin staining (Alexa Fluor 568-Phalloidin) (not shown) was used to delineate the cell outline. Bar, 10 µm.

**Figure S4.**
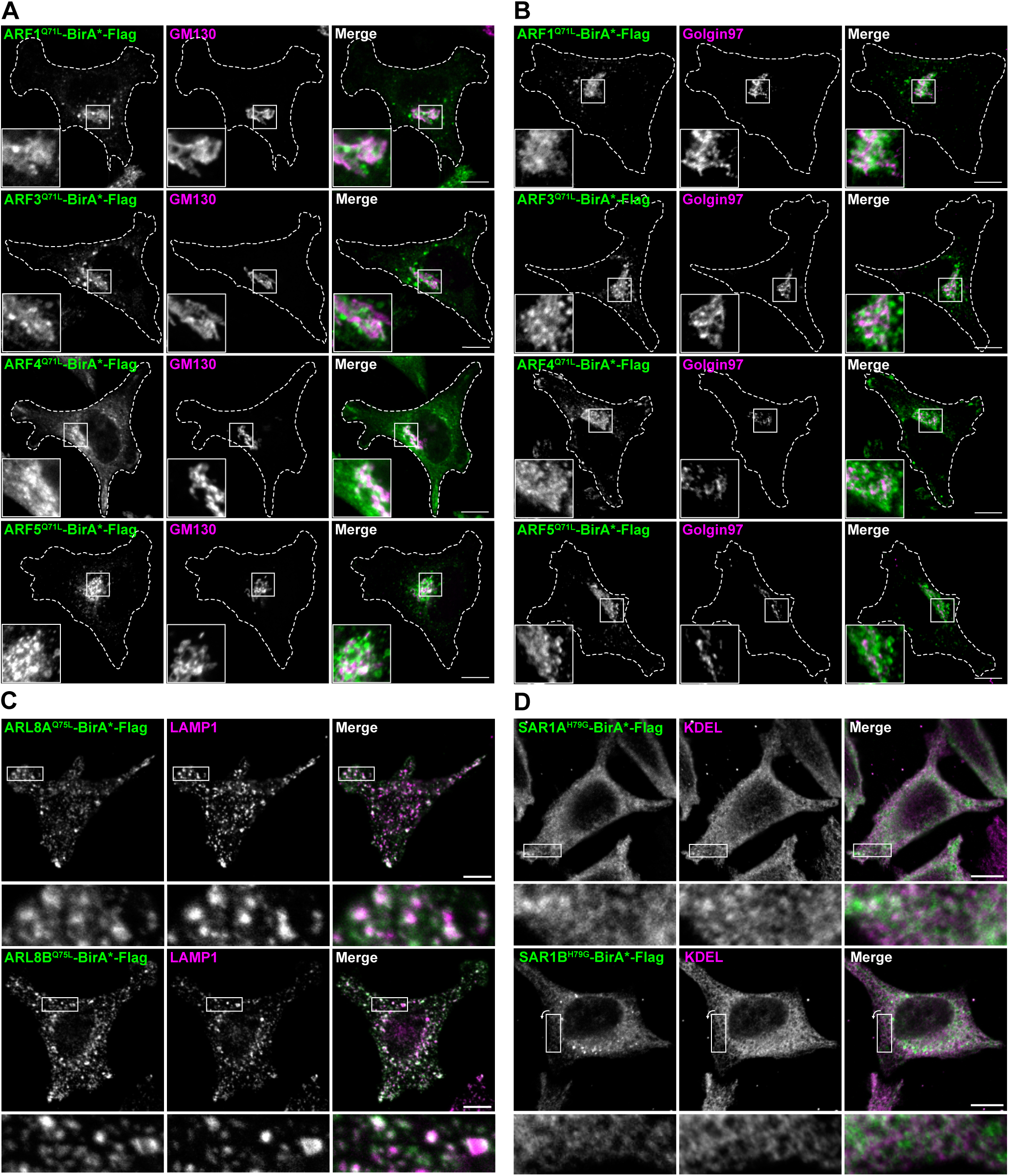
Validation of the localization of well-established ARF family members. Confocal images of co-immunostaining using anti-Flag and **(A)** anti-GM130 (*cis*-Golgi marker), **(B)** anti-Golgin97 (*trans*-Golgi marker), **(C)** anti-LAMP1 (lysosomes marker) or **(D)** anti-KDEL (the ER lumen marker). FlpIn T-REx HeLa cells were fixed and stained 24 hours after induction expression of the indicated ARF-BirA*-Flag constructs with tetracycline. Insets represent ∼2.5x magnification (A-B) and ∼4.4 magnification (C-D). Bars, 10 µm.

**Figure S5.**
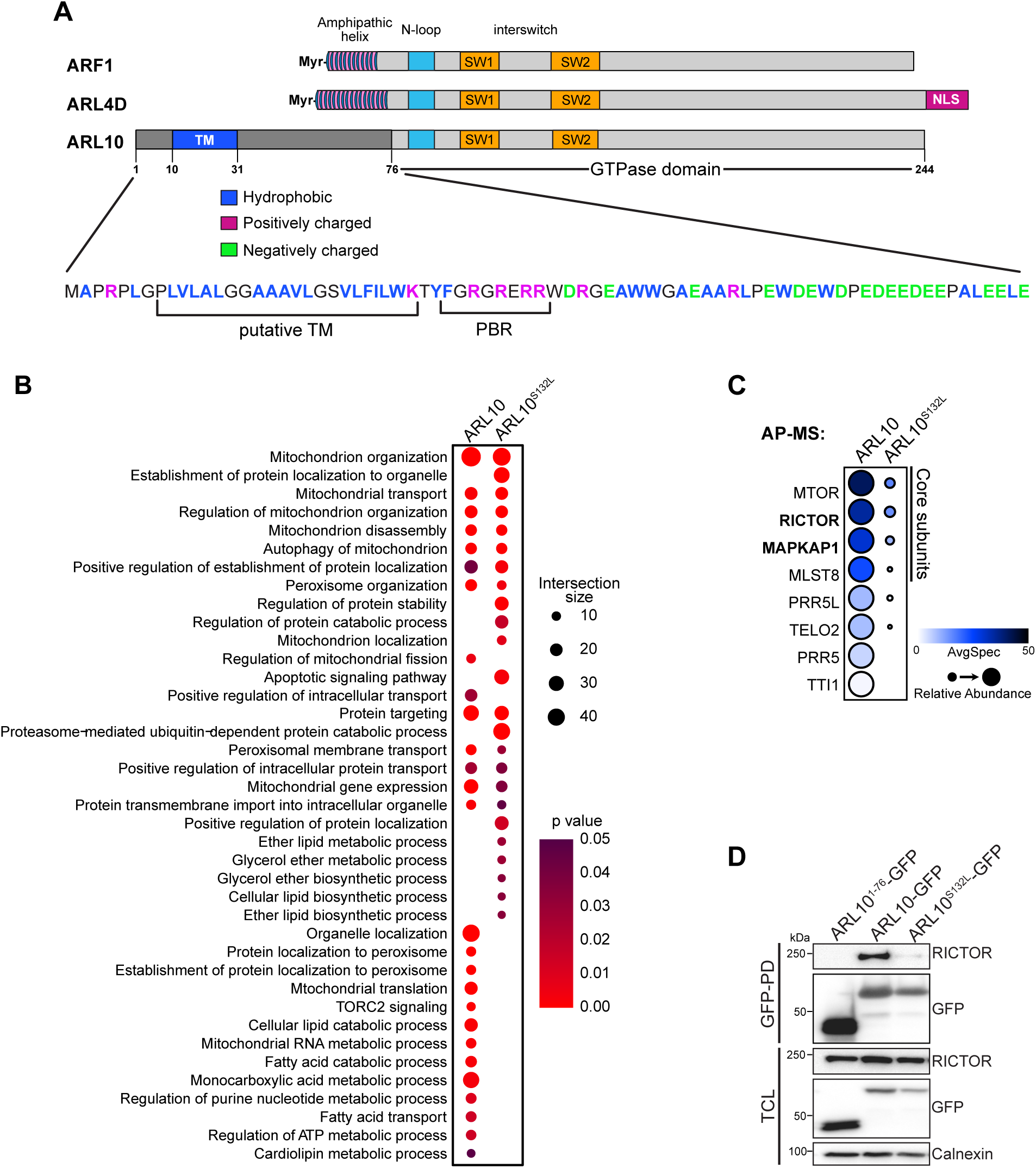
ARL10 is involved in TORC2 signaling and interacts with RICTOR. **(A)** Schematic representation of ARL4D and ARL10 GTPase features as compared to ARF1. Both ARL4D and ARL10 contain the core ARF GTPase region (in light grey) that includes the N-loop, switch 1 and switch 2 regions. ARL4D contains a nuclear localization signal (NLS) in C-terminus. The Myristoylated (Myr-) amphipathic helix is conserved in ARL4D but is replaced in ARL10 by a 76 amino acid sequence containing a predicted transmembrane (TM) domain enriched in hydrophobic residues followed by a stretch of positively charged (polybasic region, PBR) amino acids region. **(B)** Overrepresentation analyses of GO biological processes enriched from the AP-MS of ARL10-GFP (WT) and constitutively active (S132L) in FlpIn T-REx HeLa cells. The intersection size represents the number of proteins associated with a biological process. **(C)** Dotplot showing AP-MS protein-protein interactions of the mTORC2 complex retrieved in ARL10-GFP (WT) and constitutively active (S132L) in FlpIn T-REx HeLa cells. Darker circle color represents higher spectral counts while circle size represents relative abundance. AvgP ≥ 0.95. **(D)** Co-immunoprecipitation of RICTOR by pulldown (PD) of ARL10-GFP WT or constitutively active in FlpIn T-REx HeLa cells. The GFP targeted to mitochondria generated by fusion of ARL101-76 to GFP was used as control. Calnexin was used as loading control in the total cell lysate (TLC). Data are representative of n=3 experiments.

**Figure S6.**
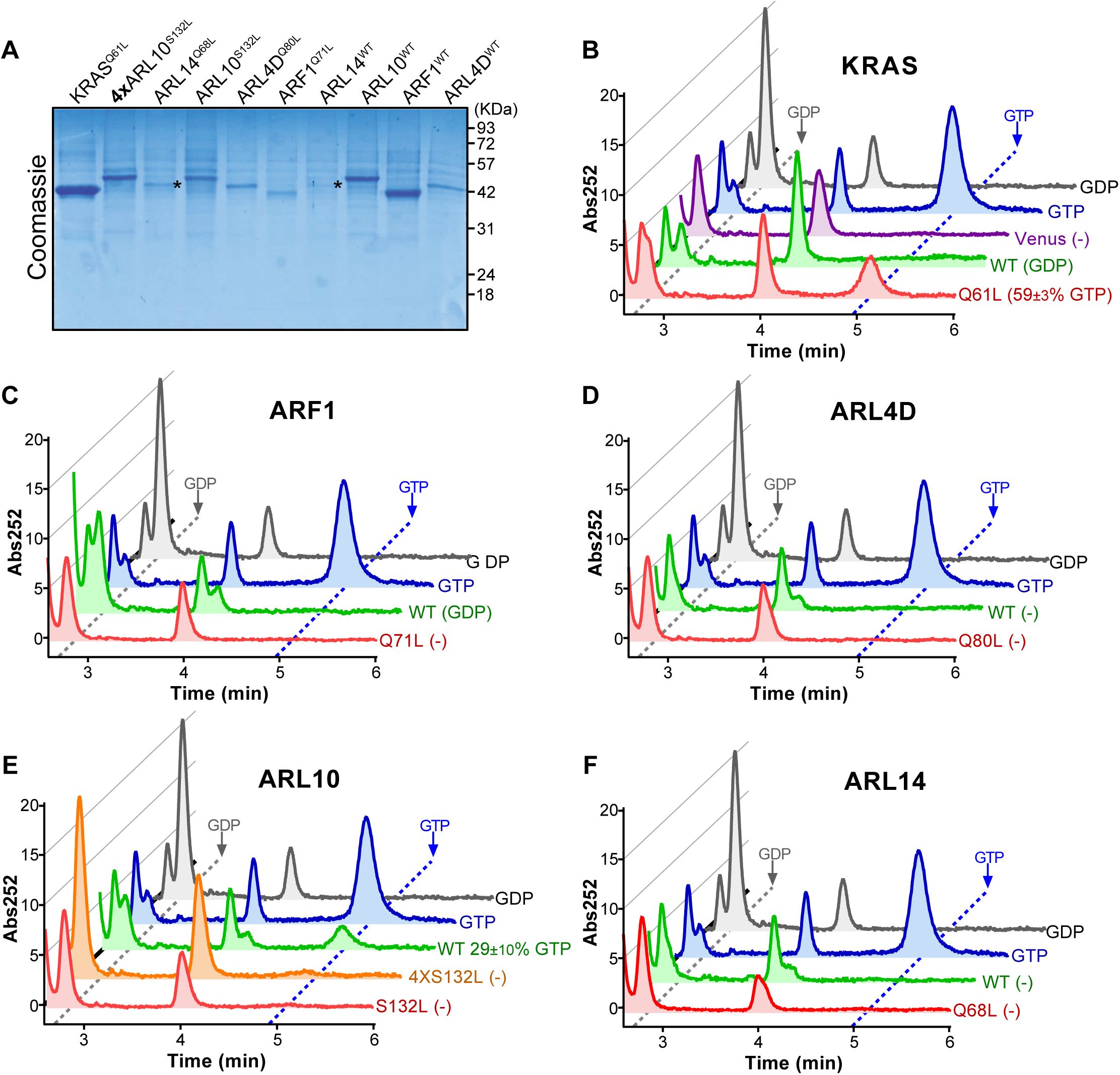
Constitutively active mutants of ARF and RAS GTPases presents differential nucleotide binding. **(A)** Coomassie stanning showing the purified fusion proteins used in (B) The asterisk (*) indicates the bands of interest for ARL14. **(B-F)** Ion-pair reversed-phase high-performance liquid chromatography performed on Venus-tagged proteins in their wild-type (in green) or constitutively active form (in red) purified after overexpression in HEK293T cells. Nucleotide alone (GDP in black and GTP in blue) and overexpression of an empty vector (Venus in purple) were used as controls. Absorbance at 252 nm was measured for 6 minutes.

**Figure S7.**
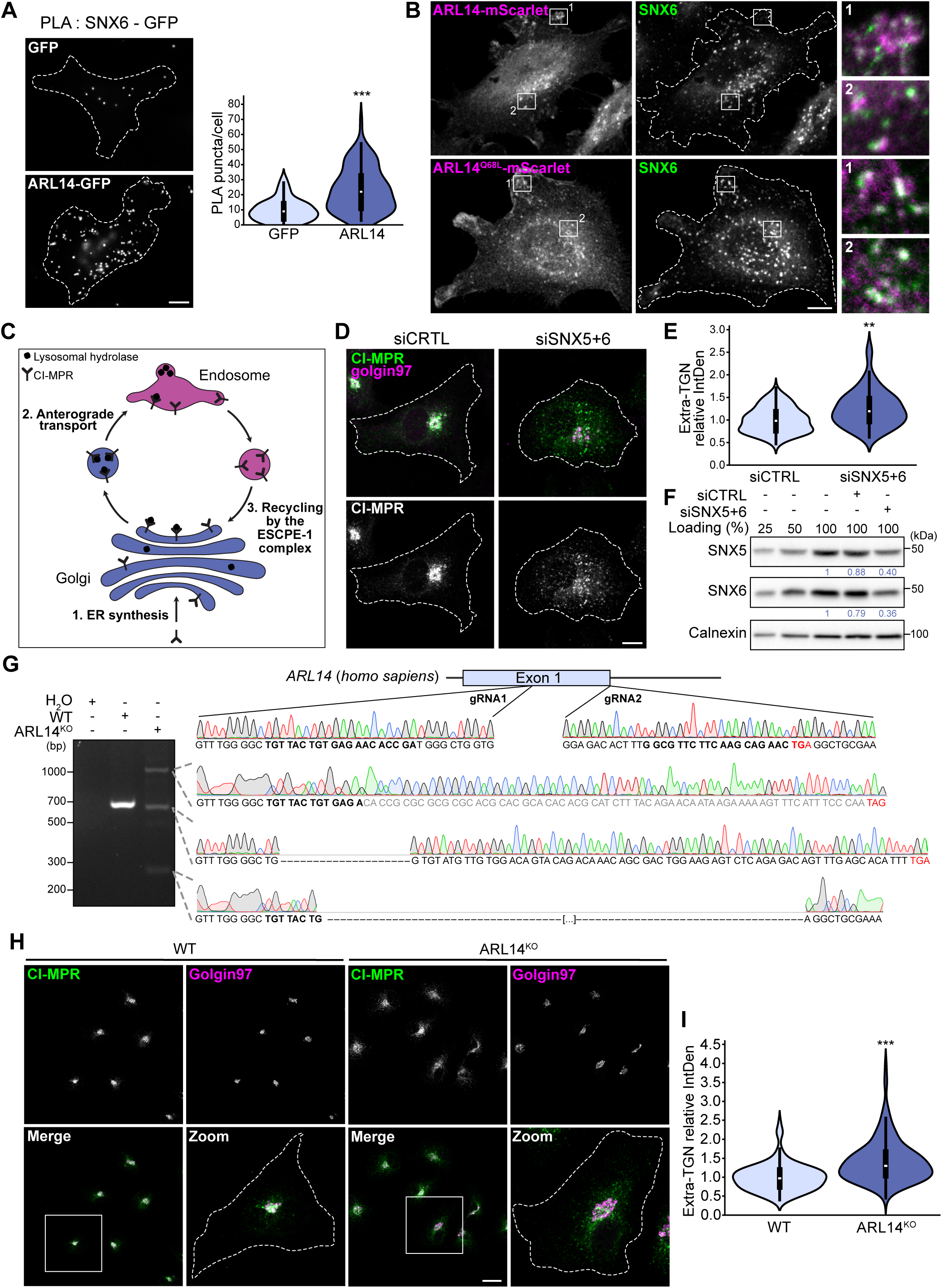
SNX6 is a close proximity interactor of ARL14. **(A)** PLA of endogenous SNX6 and overexpressed ARL14 WT tagged with GFP performed in FlpIn T-REx HeLa cells. Cells expressing GFP alone were used as a control. Bar, 10 µm. Violin plot showing the quantification of the number of PLA puncta per cell for each condition. Data are representative of n=3 independent experiments (total of 90 cells per condition). *P* value was calculated by Student *t*-test; \*\*\**P*< 0.0001. **(B)** SNX6 immunostaining of FlpIn T-REx HeLa expressing ARL14-mScarlet WT or constitutively active (Q68L). Merged images represents a ∼4.2x magnification. Bar, 10 µm. **(C)** Schematic representation of CI-MPR trafficking. **(D)** Co-immunostainings using anti-Golgin97 as TGN marker and anti-CI-MPR of HeLa cells transfected with siCTRL or siSNX5 and siSNX6. Bar, 10µm. **(E)** Violin plot showing the quantification of the integrated density (IntDen) outside the TGN relative to the control for each condition. Data are representative of n=3 independent experiments (total of 90 cells per condition). *P* value was calculated by Student *t*-test; \*\**P*< 0.001. **(F)** Western blot showing SNX5 and SNX6 knock-down after 72h of treatment with the indicated siRNAs. Calnexin was used as loading control. Data are representative of n=3 independent experiments and quantification are shown under SNX blots. **(G)** Sequencing of the *ARL14* knock-out clone in HeLa cells. Sequences for the two gRNAs used are represented in bold and the STOP codons are identified in red. Hela cells have three copies of this region as shown by the three bands in the agarose gel. The highest band shows an insertion of a small sequence that led to a STOP codon addition. The middle band represents the generation of a single cut which shifted the reading frame and led to a STOP codon addition. The lowest band represents two cuts that led to the deletion of the region between gRNA 1 and 2. **(H)** Co-immunostainings using anti-Golgin97 as TGN marker and anti-CI-MPR in HeLa WT and ARL14 knock-out cells. Dotted lines represent cell outline delimited using F-actin staining (not shown). Zoomed in images represents a ∼2.8x magnification. Bar, 10µm. **(I)** Violin plot showing the quantification of the integrated density (IntDen) outside the TGN relative to the control for each condition presented in (H). Data are representative of n=3 independent experiments (total of 120 cells per condition). *P* value was calculated by Student *t*-test; \*\*\**P*< 0.0001.

