## Supplemental Tables 1-5 for "Mapping the global interactome of the ARF family reveals spatial organization in cellular signaling pathways"

Table S1. List of baits used for BioID experiments

| | Bait | Bait Mutation | NCBI Reference Sequence or Sequence |
|---|---|---|---|
| Classical | ARF1 | Q71L | NM_001024227 |
| | ARF1 | WT | NM_001024227 |
| | ARF3 | Q71L | NM_001659 |
| | ARF4 | Q71L | NM_001660 |
| | ARF4 | WT | NM_001660 |
| | ARF5 | Q71L | NM_001662 |
| | ARF6 | Q67L | NM_001663 |
| | ARF6 | WT | NM_001663 |
| ARF-Like | ARL1 | Q71L | NM_001177 |
| | ARFRP1 | Q79L | NM_003224 |
| | ARL2 | Q70L | NM_001667 |
| | ARL3 | Q71L | NM_004311 |
| | ARL4A | Q79L | NM_005738 |
| | ARL4C | Q72L | NM_005737 |
| | ARL4D | Q80L | NM_001661 |
| | ARL5A | Q70L | NM_012097 |
| | ARL5B | Q70L | NM_178815 |
| | ARL5C | P70L | NM_001143968 |
| | ARL6 | Q73L | NM_001323513.2 |
| | ARL8A | Q75L | NM_138795 |
| | ARL8B | Q75L | NM_018184 |
| | ARL9 | S151L | NM_001363794.2 |
| | ARL10 | S132L | NM_173664 |
| | ARL11 | Q67L | NM_138450 |
| | ARL13A | D75L | NM_001162490 |
| | ARL13B | G75L | NM_182896 |
| | ARL14 | Q68L | NM_025047 |
| | ARL15 | A86L | NM_019087 |
| | ARL16 | WT | NM_001040025.3 |
| SAR | SAR1A | H79G | NM_001142648 |
| | SAR1A | WT | NM_001142648 |
| | SAR1B | H79G | NM_001033503 |
| Control | Empty vector | NA | NA |
| | EGFP | NA | ATGGTGAGCAAGGGCGAGGAGCTGTTCACCGGGGTGGTGCCCATCCTGGTC<br>GAGCTGGACGGCGACGTAAACGGCCACAAGTTCAGCGTGTCCGGCGAGGG<br>CGAGGGCGATGCCACCTACGGCAAGCTGACCCTGAAGTTCATCTGCACCAC<br>CGGCAAGCTGCCCGTGCCCTGGCCCACCCTCGTGACCACCCTGACCTACG<br>GCGTGCAGTGCTTCAGCCGCTACCCCGACCACATGAAGCAGCACGACTTCT<br>TCAAGTCCGCCATGCCCGAAGGCTACGTCCAGGAGCGCACCATCTTCTTCA<br>AGGACGACGGCAACTACAAGACCCGCGCCGAGGTGAAGTTCGAGGGCGAC<br>ACCCTGGTGAACCGCATCGAGCTGAAGGGCATCGACTTCAAGGAGGACGGC<br>AACATCCTGGGGCACAAGCTGGAGTACAACTACAACAGCCACAACGTCTAT<br>ATCATGGCCGACAAGCAGAAGAACGGCATCAAGGTGAACTTCAAGATCCGC<br>CACAACATCGAGGACGGCAGCGTGCAGCTCGCCGACCACTACCAGCAGAA<br>CACCCCCATCGGCGACGGCCCCGTGCTGCTGCCCGACAACCACTACCTGA<br>GCACCCAGTCCGCCCTGAGCAAAGACCCCAACGAGAAGCGCGATCACATG<br>GTCCTGCTGGAGTTCGTGACCGCCGCCGGGATCACTCTCGGCATGGACGAG<br>CTGTACAAGTAA |
| | EGFP-CAAX | NA | ATGGTGAGCAAGGGCGAGGAGCTGTTCACCGGGGTGGTGCCCATCCTGGTC<br>GAGCTGGACGGCGACGTAAACGGCCACAAGTTCAGCGTGTCCGGCGAGGG<br>CGAGGGCGATGCCACCTACGGCAAGCTGACCCTGAAGTTCATCTGCACCAC<br>CGGCAAGCTGCCCGTGCCCTGGCCCACCCTCGTGACCACCCTGACCTACG<br>GCGTGCAGTGCTTCAGCCGCTACCCCGACCACATGAAGCAGCACGACTTCT<br>TCAAGTCCGCCATGCCCGAAGGCTACGTCCAGGAGCGCACCATCTTCTTCA<br>AGGACGACGGCAACTACAAGACCCGCGCCGAGGTGAAGTTCGAGGGCGAC<br>ACCCTGGTGAACCGCATCGAGCTGAAGGGCATCGACTTCAAGGAGGACGGC<br>AACATCCTGGGGCACAAGCTGGAGTACAACTACAACAGCCACAACGTCTAT<br>ATCATGGCCGACAAGCAGAAGAACGGCATCAAGGTGAACTTCAAGATCCGC<br>CACAACATCGAGGACGGCAGCGTGCAGCTCGCCGACCACTACCAGCAGAA<br>CACCCCCATCGGCGACGGCCCCGTGCTGCTGCCCGACAACCACTACCTGA<br>GCACCCAGTCCGCCCTGAGCAAAGACCCCAACGAGAAGCGCGATCACATG<br>GTCCTGCTGGAGTTCGTGACCGCCGCCGGGATCACTCTCGGCATGGACGAG<br>CTGTACAAGAAGAAGAAAAAGAAGTCAAAGACAAAGTGTGTAATTATGTAG |

**Table S2. List of antibodies used in this study**

| Primary Antibodies for WB | Source | Cat# | Dilution | Incubation |
|---|---|---|---|---|
| Mouse anti-calnexin (E10) | Santa Cruz Biotechnology | sc-46669 | 1:1000 | overnight 4°C |
| anti-Flag-HRP | Sigma-Aldrich | A8592 | 1:8000 | 1h room temp |
| rabbit anti-GAPDH | Santa Cruz Biotechnology | SC-25778 | 1:5000 | 1h room temp |
| Rabbit anti-GFP | Thermo Fisher Scientific | A-11122 | 1:2000 | overnight 4°C |
| Rabbit anti-Rictor (D16H9) | Cell Signaling Technology | 9476S | 1:1000 | overnight 4°C |
| Rabbit anti-SNX5 | AbCam | ab180520 | 1:500 | overnight 4°C |
| Mouse anti-SNX6 | Santa Cruz Biotechnology | sc-365965 | 1:1000 | overnight 4°C |
| Rabbit anti-M6PR | AbCam | ab124767 | 1:10 000 | overnight 4°C |
| Rabbit anti-Golgin 97 | Proteintech | 12640-1-AP | 1:2000 | overnight 4°C |
| anti-Streptavidin-HRP | BD Biosciences | 554066 | 1:25000 | 1h room temp |

| Primary Antibodies for IF and PLA | Source | Cat# | Dilution | Incubation | Application |
|---|---|---|---|---|---|
| Rabbit anti-DYKDDDDK Tag (D6W5B) | Cell Signaling Technology | 14793S | 1:1000 | 1h 37°C | IF |
| Mouse anti-Golgin 97 (CDFX) | Santa Cruz Biotechnology | sc-59820 | 1:000 | 1h 37°C | IF |
| Rabbit anti-PLD1 | Cell Signaling Technology | 3832S | 1:100 | 1h 37°C | IF |
| Mouse anti-SNX1 | BD Biosciences | 611482 | 1:100 | 1h 37°C | IF |
| Mouse anti-SNX6 | Santa Cruz Biotechnology | sc-365965 | 1:100 | 1h 37°C | IF |
| Mouse anti-TOM20 | Proteintech | 66777-1-Ig | 1:500 | 1h 37°C | IF |
| Rabbit anti-PEX14 | Sigma-Aldrich | HPA049231 | 1:150 | 1h 37°C | IF |
| Rabbit anti-GM130 | Proteintech | 11308-1-AP | 1:200 | 1h 37°C | IF |
| Rabbit anti-M6PR | AbCam | ab124767 | 1:1000 | 1h 37°C | IF |
| Mouse anti-HA-tag | Cell Signaling Technology | 2367S | 1:500 | 1h 37°C | PLA |
| Rabbit anti-GFP | Thermo Fisher Scientific | A-11122 | 84:20:00 | overnight 4°C | PLA |

| Secondary Antibodies | Source | Cat# | Dilution | Incubation | Application |
|---|---|---|---|---|---|
| Secondary IgG-HRP anti-mouse | Sigma-Aldrich | A4416 | 1:5000 | 1h room temp | WB |
| Secondary IgG-HRP anti-rabbit | Santa Cruz | SC-2357 | 1:5000 | 1h room temp | WB |
| Alexa Fluor 488 chicken anti-rabbit IgG | Thermo Fisher Scientific | A21441 | 1:500 | 30min room temp | IF |
| Alexa Fluor 488 chicken anti-mouse IgG | Thermo Fisher Scientific | A21200 | 1:500 | 30min room temp | IF |
| Alexa Fluor 568 goat anti-rabbit IgG | Thermo Fisher Scientific | A11011 | 1:500 | 30min room temp | IF |
| Alexa Fluor 568 goat anti-mouse IgG | Thermo Fisher Scientific | A11031 | 1:500 | 30min room temp | IF |
| Alexa Fluor 633 goat anti-mouse IgG | Thermo Fisher Scientific | A21050 | 1:500 | 30min room temp | IF |

| Staining reagents | Source | Cat# | Dilution | Incubation |
|---|---|---|---|---|
| Alexa Fluor 568 Phalloidin | Thermo Fisher | A22284 | 1:1000 | 1h room temp |
| Alexa Fluor 647 Streptavidin | Thermo Fisher | S32357 | 1:1000 | 1h room temp |

**Table S3. List of plasmids used in this study**

| Plasmid | Source/Ref. | Application |
|---|---|---|
| pcDNA5-pDEST-FRT-BirA*-Flag-CT | A.C.Gingras, Lunenfeld Tanenbaum Research Institute (Couzens et al., 2013) | BioID, WB |
| pcDNA5-pDEST-FRT-ARF1Q71L-BirA*-Flag | J.F. Coté, IRCM (Patel et al., 2011) | BioID, IF, WB |
| pcDNA5-pDEST-FRT-ARF3$^{Q71L}$-BirA*-Flag | J.F. Coté, IRCM (Patel et al., 2011) | BioID, IF, WB |
| pcDNA5-pDEST-FRT-ARF4$^{Q71L}$-BirA*-Flag | J.F. Coté, IRCM (Patel et al., 2011) | BioID, IF, WB |
| pcDNA5-pDEST-FRT-ARF5$^{Q71L}$-BirA*-Flag | J.F. Coté, IRCM (Patel et al., 2011) | BioID, IF, WB |
| pcDNA5-pDEST-FRT-ARF6$^{Q67L}$-BirA*-Flag | J.F. Coté, IRCM (Patel et al., 2011) | BioID, IF, WB |
| pcDNA5-pDEST-FRT-ARL1$^{Q71L}$-BirA*-Flag | J.F. Coté, IRCM (Patel et al., 2011) | BioID, IF, WB |
| pcDNA5-pDEST-FRT-ARFRP1$^{Q79L}$-BirA*-Flag | J.F. Coté, IRCM (Patel et al., 2011) | BioID, IF, WB |
| pcDNA5-pDEST-FRT-ARL2$^{Q70L}$-BirA*-Flag | J.F. Coté, IRCM (Patel et al., 2011) | BioID, IF, WB |
| pcDNA5-pDEST-FRT-ARL3$^{Q71L}$-BirA*-Flag | J.F. Coté, IRCM (Patel et al., 2011) | BioID, IF, WB |
| pcDNA5-pDEST-FRT-ARL4a$^{Q79L}$-BirA*-Flag | J.F. Coté, IRCM (Patel et al., 2011) | BioID, IF, WB |
| pcDNA5-pDEST-FRT-ARL4c$^{Q72L}$-BirA*-Flag | J.F. Coté, IRCM (Patel et al., 2011) | BioID, IF, WB |
| pcDNA5-pDEST-FRT-ARL4d$^{Q80L}$-BirA*-Flag | J.F. Coté, IRCM (Patel et al., 2011) | BioID, IF, WB |
| pcDNA5-pDEST-FRT-ARL5a$^{Q70L}$-BirA*-Flag | J.F. Coté, IRCM (Patel et al., 2011) | BioID, IF, WB |
| pcDNA5-pDEST-FRT-ARL5b$^{Q70L}$-BirA*-Flag | J.F. Coté, IRCM (Patel et al., 2011) | BioID, IF, WB |
| pcDNA5-pDEST-FRT-ARL5c$^{P70L}$-BirA*-Flag | J.F. Coté, IRCM (Patel et al., 2011) | BioID, IF, WB |
| pcDNA5-pDEST-FRT-ARL6$^{Q73L}$-BirA*-Flag | This study | BioID, IF, WB |
| pcDNA5-pDEST-FRT-ARL8a$^{Q75L}$-BirA*-Flag | J.F. Coté, IRCM (Patel et al., 2011) | BioID, IF, WB |
| pcDNA5-pDEST-FRT-ARL8b$^{Q75L}$-BirA*-Flag | J.F. Coté, IRCM (Patel et al., 2011) | BioID, IF, WB |
| pcDNA5-pDEST-FRT-ARL9$^{S151L}$-BirA*-Flag | This study | BioID, IF, WB |
| pcDNA5-pDEST-FRT-ARL10$^{S132L}$-BirA*-Flag | J.F. Coté, IRCM (Patel et al., 2011) | BioID, IF, WB |
| pcDNA5-pDEST-FRT-ARL11$^{Q67L}$-BirA*-Flag | J.F. Coté, IRCM (Patel et al., 2011) | BioID, IF, WB |
| pcDNA5-pDEST-FRT-ARL13a$^{D75L}$-BirA*-Flag | J.F. Coté, IRCM (Patel et al., 2011) | BioID, IF, WB |
| pcDNA5-pDEST-FRT-ARL13b$^{G75L}$-BirA*-Flag | J.F. Coté, IRCM (Patel et al., 2011) | BioID, IF, WB |
| pcDNA5-pDEST-FRT-ARL14$^{Q68L}$-BirA*-Flag | J.F. Coté, IRCM (Patel et al., 2011) | BioID, IF, WB |
| pcDNA5-pDEST-FRT-ARL15$^{A86L}$-BirA*-Flag | J.F. Coté, IRCM (Patel et al., 2011) | BioID, IF, WB |
| pcDNA5-pDEST-FRT-ARL15$^{A86L}$-BirA*-Flag | J.F. Coté, IRCM (Patel et al., 2011) | BioID, IF, WB |
| pcDNA5-pDEST-FRT-ARL16-BirA*-Flag | This study | BioID, IF, WB |
| pcDNA5-pDEST-FRT-SAR1a$^{H79G}$-BirA*-Flag | J.F. Coté, IRCM (Patel et al., 2011) | BioID, IF, WB |
| pcDNA5-pDEST-FRT-SAR1b$^{H79G}$-BirA*-Flag | J.F. Coté, IRCM (Patel et al., 2011) | BioID, IF, WB |
| pcDNA5-pDEST-FRT-TRIM23$^{458I}$-BirA*-Flag | J.F. Coté, IRCM (Patel et al., 2011) | BioID, IF, WB |
| pcDNA5-pDEST-BirA*-Flag-EGFP | J.F. Coté, IRCM (Bagci et al., 2020) | BioID, WB |
| pcDNA5-pDEST-BirA*-FLAG-EGFP-CAAX | J.F. Coté, IRCM (Bagci et al., 2020) | BioID, WB |
| pcDNA5-pDEST-FRT-ARL14-EGFP | This study | IF, PLA |
| pcDNA5-pDEST-FRT-ARL14$^{S27N}$-EGFP | This study | IF, PLA |
| pcDNA5-pDEST-FRT-ARL14$^{Q68L}$-EGFP | This study | IF, PLA |
| pcDNA5-pDEST-FRT-ARL14-mScarlet | This study | IMPACT |
| pcDNA5-pDEST-FRT-ARL14$^{S27N}$-mScarlet | This study | IMPACT |
| pcDNA5-pDEST-FRT-ARL14$^{Q68L}$-mScarlet | This study | IMPACT |
| pcDNA5-pDEST-FRT-ARL10-EGFP | This study | IF, co-IP, AP-MS |
| pcDNA5-pDEST-FRT-ARL10S91N-EGFP | This study | IF, co-IP, AP-MS |
| pcDNA5-pDEST-FRT-ARL10$^{S132L}$-EGFP | This study | IF, co-IP, AP-MS |
| pCGN-hPLD1b | M.A. Frohman, Stony Brook Cancer Center (Hammond et al., 1995) | PLA |
| pOG44 | Thermo Fisher Scientific (12536017) | Transfections |
| pDONR221 | Thermo Fisher Scientific (V600520) | Cloning |
| pcDNA5-pDEST-FRT-mScarlet-CT | This study | Cloning |
| pcDNA5-pDEST-FRT-EGFP-CT | A.C.Gingras, Lunenfeld Tanenbaum Research Institute (Kean et al., 2011) | Cloning |
| YFP-hPOP1-C1 | This study | Transfections |
| pGEX-4T-1-ARL14 | This study | NMR, ITC |
| pGEX-4T-1-ARL11 | This study | NMR, ITC |
| pDEST-pcDNA-ARF1-Venus | This study | HPLC |
| pDEST-pcDNA-ARF1$^{Q71L}$-Venus | This study | HPLC |
| pDEST-pcDNA-ARL14-Venus | This study | HPLC |
| pDEST-pcDNA-ARL14$^{Q68L}$-Venus | This study | HPLC |
| pDEST-pcDNA-ARL4D-Venus | This study | HPLC |
| pDEST-pcDNA-ARL4D$^{Q80L}$-Venus | This study | HPLC |
| pDEST-pcDNA-ARL10-Venus | This study | HPLC |
| pDEST-pcDNA-ARL10$^{S132L}$-Venus | This study | HPLC |
| pDEST-pcDNA-KRAS$^{Q61L}$-Venus | This study | HPLC |
| pDEST-pcDNA-Venus | This study | HPLC |

**Table S4. List of siRNA used in this study**

| siRNA | Cat# | Sequence |
|---|---|---|
| SNX5 | siGENOME Human SNX5 (27131) siRNA - SMART pool M-012524-00-0005 | UAACAGAGCUCCUCCGAUA |
| | | GAGCAAAGACGUCAAGUUA |
| | | CAAACAAAGCUCUGGAUAA |
| | | CUACGAAGCCCGACUUUGA |
| SNX6 | siGENOME Human SNX6 (58533) siRNA - SMART pool M-017557-01-0005 | GAUGAAGACCUCAAACUUU |
| | | UAAAUCAGCAGAUGGAGUA |
| | | CAAGAAGAGUUGCUGCAUU |
| | | ACUUAGUAGUUAUGCGAGU |
| siCTRL | siGENOME Non-Targeting siRNA #1 - D-001210-01-20 | UAGCGACUAAACACAUCAA |

**Table 5. List of oligos used in this study**

| PCR | Target | Application | Forward | Reverse |
|---|---|---|---|---|
| | mScarlet | KpnI/mScarlet/XhoI PCR amplification | 5'-AAGCTTGGTACCATGGTGAGCAAGGGC-3' | 5'-CATCATCTCGAGTTACTTGTACAGCTCGTCCATGCC-3' |
| | ARL10-GFP | Create ARL10 a.a 1-31-GFP | 5'-ATGGTGAGCAAGGGCGAGGA-3' | 5'-GTAGGTCTTCCAGAGGATGAAGAGCACCG-3' |
| | ARL10-GFP | Create ARL10 a.a 1-76-GFP | 5'-ATGGTGAGCAAGGGCGAGGA-3' | 5'-TTCCAGCTCCTCCAGCGCC-3' |
| | ARL10-GFP | Create ARL10 Δ7-31_GFP | 5'-GGCCGCGGCCGAGAG-3' | 5'-GAACAGCGGCCGCGG-3' |
| | ARL10-GFP | Create ARL10 Δ32-76_GFP | 5'-CGCGAGGTGCTGGTGC-3' | 5'-CTGGTAGGTCTTCCAGAGGATGAAGAGC-3' |
| | ARL10-GFP | Create ARL10 Δ7-76_GFP | 5'-CGCGAGGTGCTGGTGC-3' | 5'-CTGCAGCGGCCGCGG-3' |
| | mARL14 | Genotyping ARL14 embryo | 5’-GCTGTGCGGTCACTGGAGAA-3’ | 5’-CCAAAAGTTTGCCGACTCAGC-3’ |
| | mARL14 | Genotyping ARL14 embryo in FLAG | 5’-GCTGTGCGGTCACTGGAGAA-3’ | 5’- GTCATGATCTTTATAATCACCGTCATGGTC-3’ |
| | hARL14 | Validation of ARL14 KO cells | 5'-TCACAGTCTGGGATGTTGGA-3' | 5'-AGTGATCATTGCCACAGTTTGC-3' |

| CRISPR | Name | Application | Sequence |
|---|---|---|---|
| | gRNA_mARL14 | To target CRISPR machinery to ARL14 | 5'-AGTTTCTGAAAAGCTACCGA-3' |
| | ssDNA_ARL14_3xFlag | Template to insert 3xFlag | 5'AATAAAACAGATTTCCCATTTCCACTTCATC ATTGTCTCACTTGTCATCGTCATCCTTGTAGT CGATGTCATGATCTTTATAATCACCGTCATGG TCTTTGTAGTCCCCTCCACTGCCACCTTTCT GCTTGAAGATTGCTAAGGTCTCTCTTGTTCTT |
| | gRNA1_hARL14_F | To target CRISPR machinery to ARL14 | 5'-CACCGTGTTACTGTGAGAACACCGA-3' |
| | gRNA1_hARL14_R | To target CRISPR machinery to ARL14 | 5'-AAACTCGGTGTTCTCACAGTAACAC-3' |
| | gRNA2_hARL14_F | To target CRISPR machinery to ARL14 | 5'-CACCGGCGTTCTTCAAGCAGAACTG-3' |
| | gRNA2_hARL14_R | To target CRISPR machinery to ARL14 | 5'-AAACCAGTTCTGCTTGAAGAACGCC-3' |
